# Human hormone seasonality

**DOI:** 10.1101/2020.02.13.947366

**Authors:** Avichai Tendler, Alon Bar, Netta Mendelsohn-Cohen, Omer Karin, Yael Korem, Lior Maimon, Tomer Milo, Moriya Raz, Avi Mayo, Amos Tanay, Uri Alon

## Abstract

Hormones control the major biological functions of stress response, growth, metabolism and reproduction. In animals these hormones show pronounced seasonality, with different set-points for different seasons. In humans, the seasonality of these hormones remains unclear, due to a lack of datasets large enough to discern common patterns and cover all hormones. Here, we analyze an Israeli health record on 46 million person-years, including millions of hormone blood tests. We find clear seasonal patterns: the effector hormones peak in winter-spring, whereas most of their upstream regulating pituitary hormones peak only months later, in summer. This delay of months is unexpected because known delays in the hormone circuits last hours. We explain the precise delays and amplitudes by proposing and testing a mechanism for the circannual clock: the gland masses grow with a timescale of months due to trophic effects of the hormones, generating a feedback circuit with a natural frequency of about a year that can entrain to the seasons. Thus, humans may show coordinated seasonal set-points with a winter-spring peak in the growth, stress, metabolism and reproduction axes.

## Introduction

The major biological functions in mammals- growth, reproduction, metabolism and stress adaptation - are controlled by dedicated hormonal axes. In each axis, signals from the hypothalamus cause secretion of specific pituitary hormones into the bloodstream. The pituitary hormones instruct a peripheral organ to secrete effector hormones with widespread effects on many tissues. For example, stress response is governed by the hypothalamic-pituitary-adrenal (HPA) axis: physiological and psychological stress signals cause the hypothalamus to induce secretion of ACTH from the pituitary, which instructs the adrenal cortex to secrete cortisol. These axes act to maintain physiological set-points. The set-points can change to adapt to different situations, a concept known as rheostsis (Mrosovsky, 1990). A major reason that organisms change their set-points is the seasons.

Animals show seasonal changes in the pituitary and effector hormones that govern seasonality in reproduction, activity, growth, pigmentation, morphology and migration (Gwinner, 2012). This adaptive physiology includes changes in body composition, organ size and function. In general, hormone seasonality is thought to be a dominant regulator of physiological and behavioral traits in animals (Jacobs and Wingfield, 2000; Dufty Jr et al., 2002). Animals even show these changes with a circannual rhythm when maintained in constant photoperiod and temperature conditions (Zucker, 2001; Lincoln et al., 2006; Gwinner, 2012). They cycle without external signals by means of an internal oscillator with a period of about, but not exactly, one year. The mechanism and physiological location of this circannual clock is a subject of current research. A key component is the *pars tuberalis* in the pituitary stalk, whose thyrotorph cells oscillate between high and low states of hormone production (MacGregor and Lincoln, 2008 ; Wood et al., 2015; Wood and Loudon, 2018). This area receives input on photoperiod from melatonin signals. Whether hormones show seasonality in humans has not been studied comprehensively by tracking many hormones in a large number of participants. Each axis has been studied separately, usually with small samples. These studies suggest that thyroid hormones and saliva cortisol show seasonal variation on the order of 10% (Santi *et al*., 2009; Persson et al., 2008). The studies are limited by considerations of circadian rhythms which affect cortisol and other hormones.

To study human hormone seasonality requires a large dataset with a comprehensive coverage of all hormones. Here we provide such as study using an Israeli medical record database with millions of blood tests. We address the circadian rhythm concern using the time of each test. We find coordinated seasonality with a winter/spring peak in effector hormones and a surprising antiphase between pituitary and effector hormones. We provide an explanation for this antiphase by showing that trophic effects of the hormones create a circuit in which the functional mass of the glands changes over the year and can entrain to yearly signals. The results support a winter-spring peak for human reproduction, metabolism, growth and stress adaptation.

## Results

### Data on millions of blood tests shows seasonality

We analyzed electronic medical record data from a large Israeli Health maintenance organization (Clalit), which includes millions of blood tests (Fig 1). The dataset includes measurements from about half of the Israeli population over 15 years (2002-2017) totaling 46 million life-years, with broad socioeconomic and ethnic representation.

**Figure 1.**
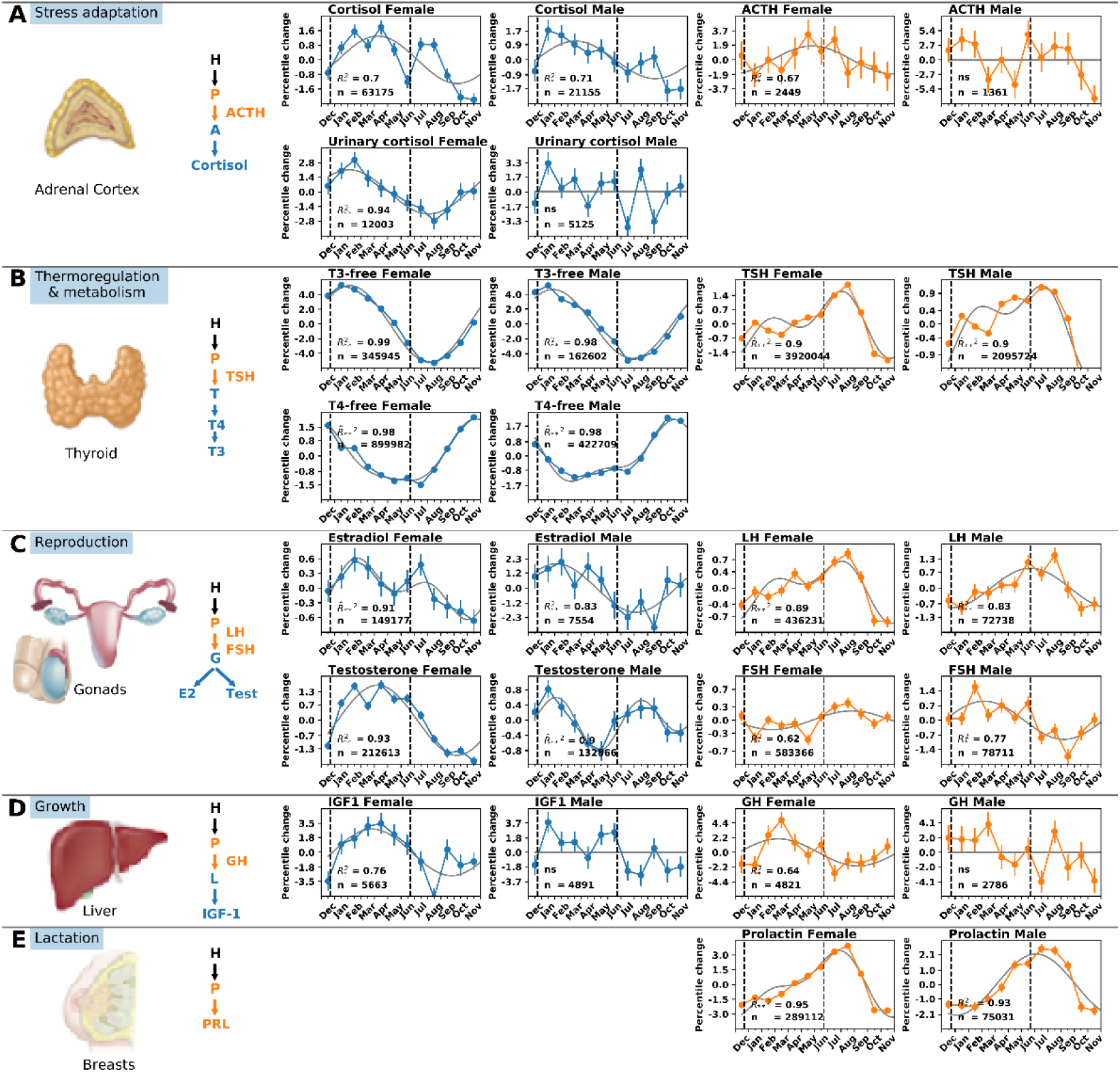
Seasonality of hypothalamic-pituitary axes hormones from Clalit medical records. (A) HPA axis with pituitary hormone ACTH and effector hormone cortisol. (B) Thyroid axis with pituitary hormone TSH (thyroid stimulating hormone) and effector hormones T4 and its derivative T3. (C) Sex axis with pituitary hormones FSH (follicular stimulating hormone) and LH (luteinizing hormone), and effector hormones testosterone and estradiol. (D) Growth axis with pituitary hormone GH (growth hormone) and effector hormone IGF1, (E) lactation pathway with pituitary hormone PRL (prolactin) that controls breast milk production. Each panel indicates the number of tests n, zero-mean cosinor model (gray line) and *R*^2^ where significant (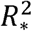 for p < 5 · 10^−2^, 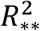 for p < 10^−3^, ns - not significant), with first or second order model selected by the Akaike criterion (second-order model is indicated by ^ above *R*). Vertical dashed lines indicate solstices Dec 21 and Jun21.

Electronic medical records have the challenge of ascertainment bias because the tests are performed for medical reasons. To address this, for each blood test we removed data from pregnant women, individuals with medical conditions that affect the blood test, and individuals that take drugs that affect the blood test (Methods).

We considered male and females separately and binned age groups from 20-80 by decades (data for age range 20-50 is similar, Fig S1). We performed quantile analysis in each decade to avoid age-related trends and outliers. Seasonality was assessed by cosinor tests (Table S1).

One concern is hormone circadian rhythms, raising the question of the time of day the test was taken relative to the person’s circadian cycle. To address this, we obtained the time of day of each test. The distribution of test times did not vary between the seasons for any of the hormones (Fig S2-S3). The seasonality conclusions presented below are unaffected by considering only tests done at specific times of day. The conclusions are also unaffected by considering only tests done at a fixed time interval (2.5h) from dawn for most tests (SI Fig S4-S5). We also compared blood tests from one of the most circadian hormones, cortisol, to urine tests that gather cortisol over 24 hours, and find similar seasonality. We conclude that we can effectively control for circadian effects (Fig S6).

The data allowed us to consider all pituitary and effector hormones in a single framework. Hormones secreted by the anterior pituitary for reproduction (LH), metabolism (TSH), stress (ACTH) and lactation (PRL), showed seasonal oscillations that peak in summer, July or August (Fig 1,2). The pituitary stress hormone ACTH in males has much fewer tests which precludes identifying seasonality. The amplitude is on the order of 1-3 percentiles. This seasonality can be detected by virtue of the large number of tests. For example, TSH exceeds 6 million blood tests, resulting in error bars smaller than the dots in Fig 1.

**Figure 2.**
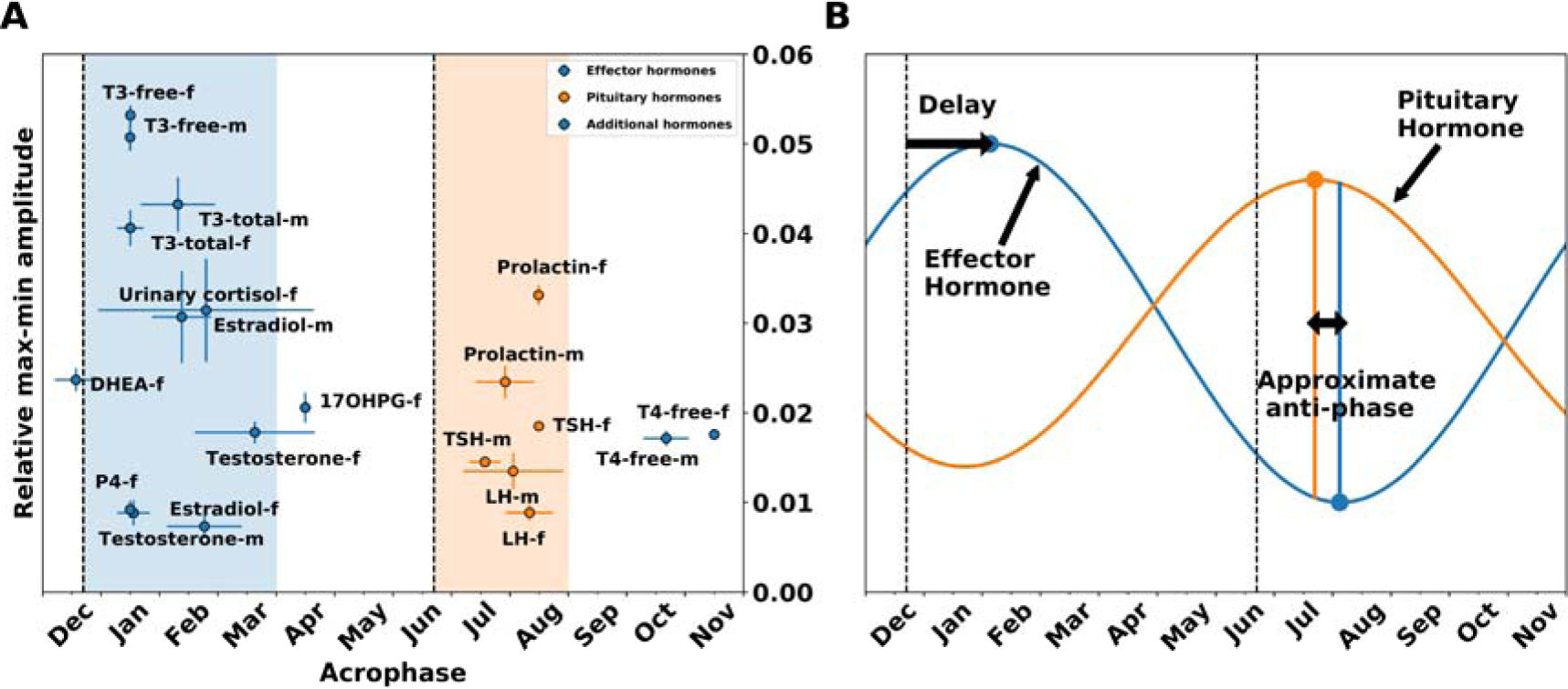
Most pituitary hormones peak in summer whereas effector hormones peak in winter-spring in Hypothalamus-Pituitary axes. (A) Peak phases (acrophase) and amplitudes of all hormones whose fit to the cosinor model exceeded R^2^ > 0.8. Dec 21 and Jun 21 are indicated (vertical dashed lines), as well as winter-April (blue region) and summer (orange region). m,f indicate male and female. (B) Schematic showing spring shift of effector hormones (blue) and approximate antiphase of the pituitary hormones (orange).

The effector hormones, secreted from peripheral organs under control of the pituitary hormones, also showed seasonality with amplitudes of 1-6 percentiles. In contrast to the summer peak of most of the pituitary hormones, the effector hormone tests peaked in the winter or spring (Fig 2). The thyroid hormone T3 peaked in winter, in agreement with previous studies (Harrop & Hopton, 1985, Maes *et al*., 1997), consistent with its role in thermogeneration. Its precursor T4 peaked in late fall. The other effector hormones peaked in late-winter or spring, including the sex hormones testosterone, estradiol, progesterone and the growth hormone IGF1. Androgen effector hormones also show similar seasonality (Fig S7). For cortisol, both blood tests and 24-hour urine tests peaked in February. This late winter peak of cortisol is in agreement with previous studies, including large studies on saliva (Persson et al., 2008) and hair (Abell *et al*., 2016) cortisol, as well as smaller studies, including a similar winter peak shifted by 6 months in the southern hemisphere (Hadlow *et al*., 2014, 2018).

Several hormones showed a secondary peak, forming a biannual rhythm. To analyze this, we used a second-order cosinor model. TSH showed a major peak in august and a minor peak in winter, in agreement with a study on 1.5 million blood tests from Italy (Santi *et al*., 2009). PRL test showed a similar biannual profile, perhaps due to the upstream regulator TRH that it shares with TSH (Friesen & Hwang, 1975). The tests for the major sex effector hormones estradiol in females and testosterone in males both showed a biannual pattern with a secondary summer peak.

There are two exceptions to the rule that pituitary hormones peak in summer. The tests for pituitary growth hormone, GH, peak in spring, close to its downstream hormone IGF1. We note that like IGF1, GH also acts as an effector hormone, because it regulates growth and metabolism, unlike many other pituitary hormones which mainly have a regulatory role and are not effector hormones. Below we offer a model that explains the GH phase based on the slow turnover of its effector organ. The second exception is FSH in males that peaks in spring.

We also considered 10 of the most common blood chemistry tests (ions, glucose, urea) (Fig 3). These tests also showed seasonal oscillations with amplitudes on the order of 0.5-8 percentiles. Their peak phases concentrated around Dec21 (shortest photoperiod) June 21 (longest photoperiod, SI). Seasonality is shown in Fig S8 and table S1.

**Figure 3.**
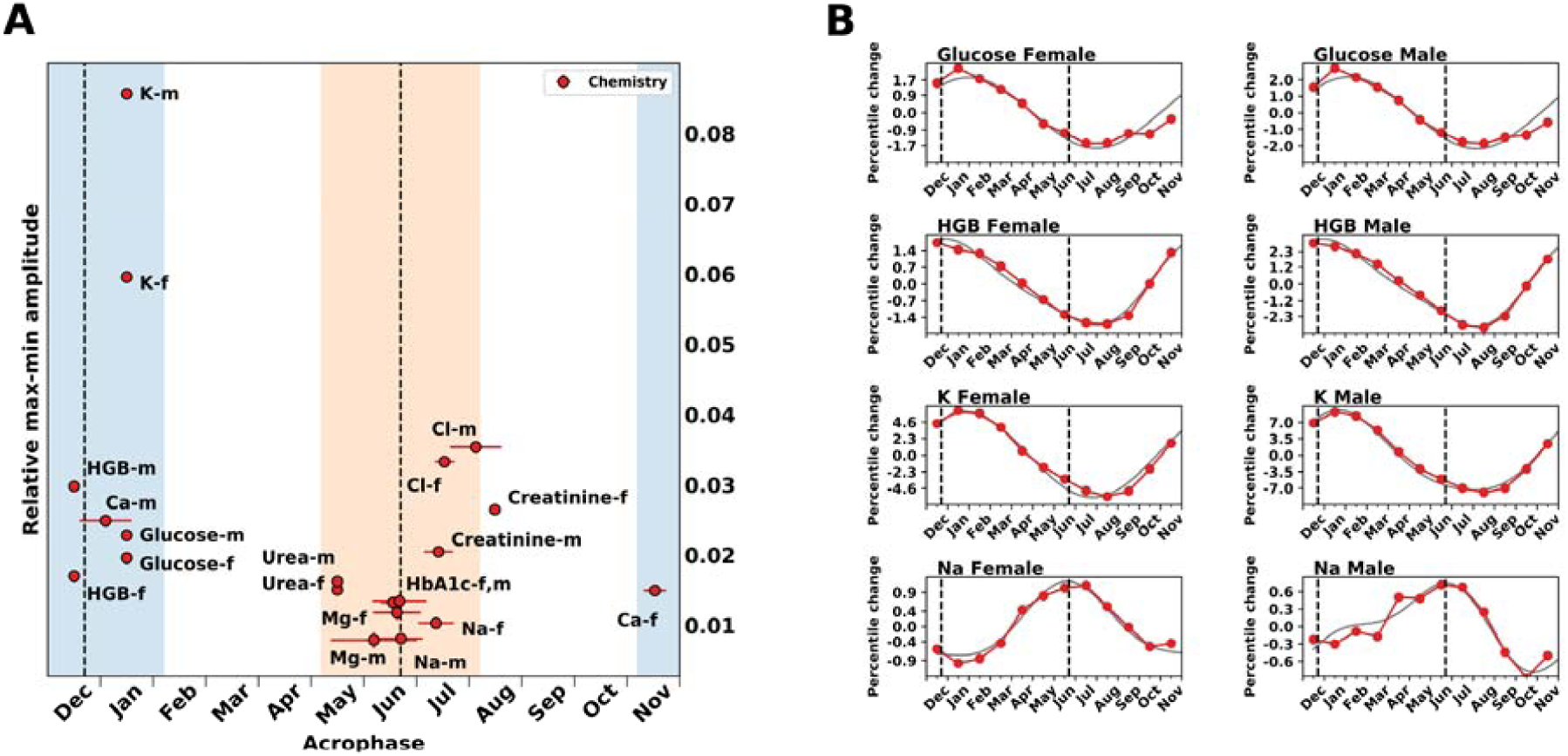
Blood chemistry tests peak around the shortest and longest photoperiods. **Amplitude and** time of peak (acrophase) of blood chemistry tests is shown, blue and orange regions are two months around the solstices Dec21 and Jun21. m,f denote male and female.

We conclude that there is a spring-delay for most of the effector hormone, shifting their peak from Dec 21 to later in the winter or to spring. Furthermore, there is a phase shift between many of the effector hormones and their upstream pituitary hormone regulators, placing them in approximate anti-phase (Fig 2B).

### Phase shifts may be due to hormone-driven changes in gland masses

The spring-delay of effector hormones and the approximate antiphase between most pituitary and effector hormones are puzzling, given the classical understanding of these axes. The classical model is that each pituitary hormone instructs the release of its effector hormones from peripheral glands and is in turn inhibited by the effector hormones by negative feedback loops (Fig 4A). Delays in these processes are on the order of minutes to days, and are negligible compared to the scale of months required to understand the spring-delay and antiphase. In contrast to the observed blood tests, the classical model predicts that pituitary hormones should coincide with their regulated hormones, and thus have the *same seasonal phase* as the effector hormones that they control (no antiphase). For hormones controlled by photoperiod, the peak should be at the time of extreme photoperiod (eg Dec21), (Fig 4A).

**Figure 4.**
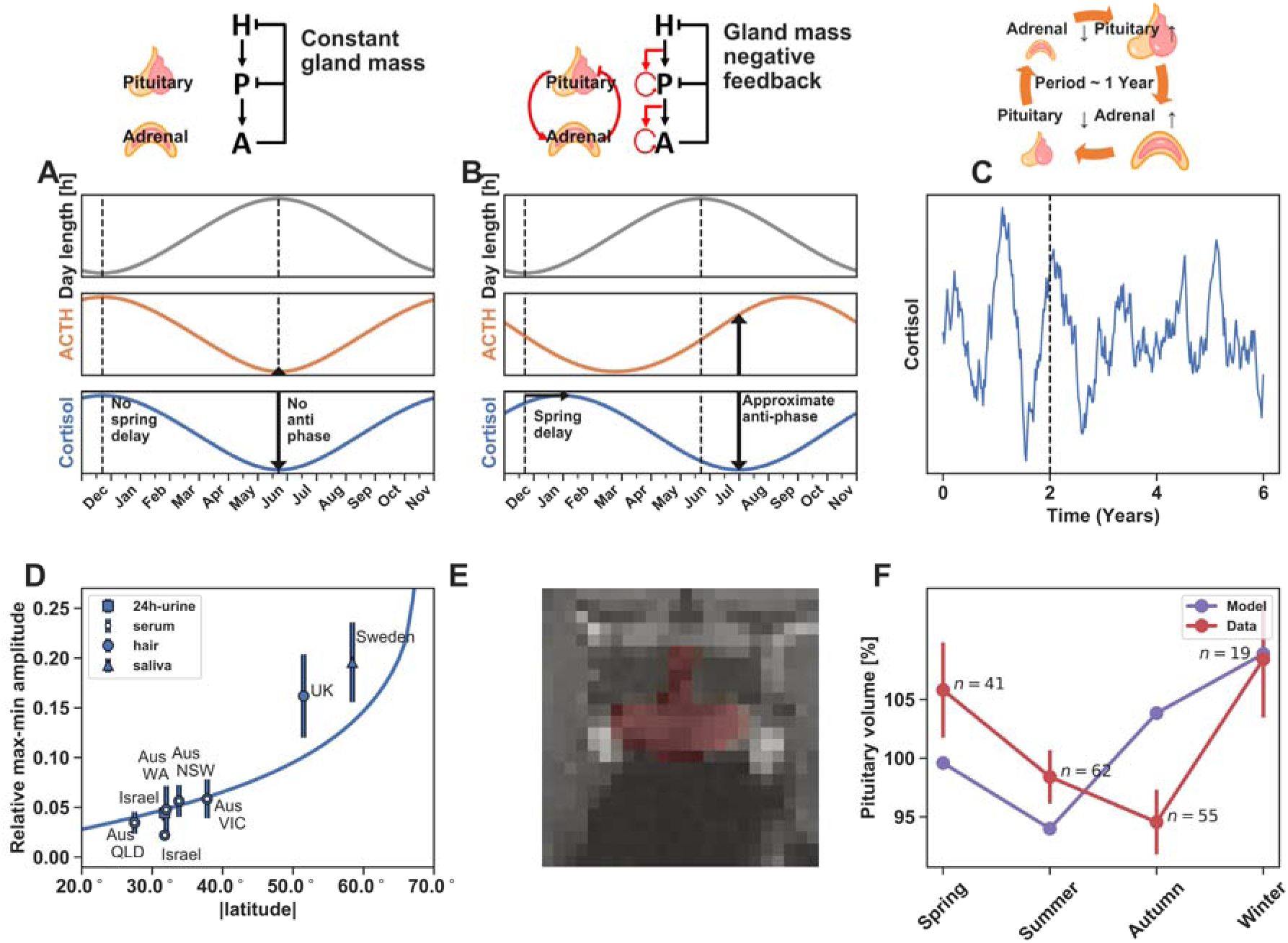
Mechanism for hormone seasonal phases based on a gland-mass oscillator. (A) Classic model of the HPA axis assumes that the mass of the cells that secrete ACTH and cortisol is constant. It predicts that an input maximal in Dec 21 will show both ACTH and cortisol peaks at Dec 21. (B) Model which considers effect of hormones as growth factors of their downstream glands (red interactions). It predicts a spring delay of cortisol and a summer peak of ACTH. (C) The gland mass model effectively generates a feedback loop in which gland masses can entrain to yearly input cycles. When provided with noise, it can oscillate even without entraining signal, as shown in stochastic simulation of two years of entrainment to a yearly photoperiod input was *u*(*t*) = 1 + *gW*(*ϕ*)*cos*(*ωt*) + *noise* followed by four years without entraining input *u*(*t*) = 1 + *noise*, where *ϕ*= 50°, *g*= 0.5, 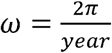. Noise was a uniformly distributed random number *u*([0.5,1.5]) that was constant over each simulated week. (D) Amplitude of cortisol seasonal variation increases with absolute latitude. Gland-mass model - blue line. Blood tests from Australia (Hadlow *et al*. 2018) (open circle), blood (open circle) and urine (square) tests from present study, hair (Abbel *et al*., 2016) (circle) from UK, and saliva (triangle) from Sweden (Persson et al., 2008). (E) Example of a pituitary segmented in an MRI image from the human connectome dataset. (F) Mean pituitary volume from MRI images binned by four seasons (red),with gland-mass model prediction (purple).

To understand the possible mechanism for the spring-delay and antiphase, we considered additional mechanisms that can provide a timescale of months. We focus on the HPA axis because it is well-studied in terms of molecular interactions (Chrousos, 1998); the other axes have analogous interactions (detailed in the SI section S8). We find that a sufficient model for the observed phase-shifts arises from adding to the classical model the effect of hormones as the primary growth factors of the tissues that they control (Fig 4B). These interactions are well-characterized, but have not been considered on the systems level. In the HPA axis, ACTH not only causes the adrenal to secrete cortisol, it also increases the growth of the adrenal cells (Kataoka, Ikehara and Hattori, 1996; Lotfi and de Mendonca 2016). Likewise, CRH not only causes corticotrophs in the pituitary to secrete ACTH, it also increases their growth rate (Westlund et al., 1985; Gertz et al., 1987; Nolan et al., 1998). Thus, the total functional mass of the corticotrophs and adrenal cortex cells are time-dependent variables. They change on the time-scale of weeks, as shown by experiments in rodents (Nolan, Thomas and Levy, 2004), due to both hyperplasia and hypertrophy. The masses of the adrenal and pituitary are also known to change in humans, growing under prolonged stress or major depression and shrinking back when the stress or depression is relieved (Parker, Schatzberg and Lyons, 2003). A similar idea, that seasonal clocks could arise from generation of tissue mass, was named by Lincoln et al the ‘cyclic histogenic hypothesis’ (Hazlerigg & Lincoln, 2011).

We modeled the functional masses together with the classic hormone circuit (eq 1-5 in methods) (Karin *et al*., 2020). On the timescale of weeks, the HPA axis acts as a damped oscillator in which the corticotroph and adrenal masses form a negative feedback loop (Fig 4C), with a typical timescale of a year (methods), that can entrain with seasonal input signals.

Simulations and analytical solutions (SI section 6) show that the model entrains to a yearly seasonal input proportional to photoperiod, maximal in Dec 21, and minimal in Jun 21, providing seasonal oscillations to the hormones. The model provides the spring delay and the approximate antiphase (Fig 4B): the effector hormone cortisol peaks in spring and the pituitary hormone ACTH peaks in late summer. The spring shift of cortisol and the antiphase between the hormones is caused by the time it takes the functional cell masses to grow and shrink.

The spring delay and antiphase are found for a wide range of model parameters: the only parameters that affect the phases are the turnover times of the cell functional masses, which can range from a week to a few months and still provide the observed antiphase to within experimental error (Fig S10). All other parameters such as hormone secretion rates and half-lives affect only the dynamics on the scale of hours, and do not measurably affect the seasonal phases.

Similar interactions exist in the other pituitary axes (Plant, 2015; Ortiga-Carvalho *et al*., 2016). Gland masses also dynamically change in these axes under control of the hormones. For example, thyroid proliferation is activated by the upstream hormone TSH, and thyroid volume changes in humans as exemplified by goiter (Dumont et al., 1992). We provide models for each axis in the SI (section S9), finding that they are sufficient to explain the phases in each axis. The models can also explain why GH peaks in spring unlike the other pituitary hormones, due to the very slow turnover time of the peripheral gland in this axis, the liver. This places GH, which has effector functions on growth and metabolism, in a similar phase to IGF1, its effector hormone. Which makes physiological sense. Due to the slow turnover of the liver, the model does not have imaginary eigenvalues. However, it can still entrain to the seasonal inputs and provide a delay from winter solstice provided by the pituitary cell turnover time.

The models can also explain the fall peak of the thyroid hormone T4, based on the architecture of cell control in the thyroid axis. The phase difference between T3 and T4 might be due to seasonality in deiodinases that converts T4 to T3 (SI section 8). Alternative putative mechanisms with a slow timescale, such as epigenetic mechanisms or seasonal parameter changes, are also considered in the SI (Fig S12).

The gland-mass models can also provide a mechanism for the circannual clock found in animals kept in constant photoperiod conditions. Although the oscillator in the model is damped in the deterministic equations, noise can cause it to show undamped oscillations at the resonance frequency of the damped oscillator (on the order of a year), Fig 4C, Fig S9. Such noise-driven oscillations have been studied in other biological systems (Alon 2019; Geva-Zatorsky et al., 2010). This model can complement models based on epigenetic (Wood et al., 2015; Wood and Loudon, 2018) and histogenic (Hazlerigg & Lincoln, 2011) mechanisms.

### Seasonality increases with latitude

The model predicts that the amplitude of seasonal variations should increase with latitude, due to greater photoperiod variation with the seasons. To test this we compared the model with studies from Australia (30°S), the UK (51°N) and Sweden (58°N) on cortisol. The amplitude of cortisol seasonality rose with absolute latitude in agreement with the model predictions (SI section 11) (Fig 4F).

### Pituitary volume shows seasonality

The functional mass model makes another testable prediction: the masses of the glands that secrete the hormones should vary with the seasons with specific phases. The total pituitary mass is made of several cell types, including somatotrophs that secrete GH, thyrotrophs that secrete TSH, corticotrophs that secrete ACTH, gonadotrophs that secrete LH/FSH and lactotrophs that secrete PRL, as well as other factors including vasculature. Since we find that most of the pituitary hormones have similar phases (Fig 1), we reasoned that one can consider total pituitary mass as a single variable. The model then predicts that the pituitary mass should peak in late spring, and thus be (perhaps surprisingly) out of phase with the levels of the pituitary hormones that peak in late summer. This antiphase is due to the inhibition by the effector hormones – a prediction that is robust to model parameters. To test this, we analyzed a dataset of MRI brain scans and computed the volume of the pituitary (SI section 12). The data and model show rough agreement (Fig 4E).

## Discussion

We find that human hormone tests show a seasonal pattern with amplitudes on the order of a few percent. Most pituitary hormones peak in late summer, and effector hormones from downstream peripheral organs peak in winter/spring.

The hormone seasonality in the present study (latitude 32°) has an amplitude on the order of a few percent. The physiological effect of such changes is not clear. Because of the coordinated peak in all axes at the same winter/spring season, and the widespread effects of each hormone on many metabolic and behavioral systems, even small changes from hormone baselines may have a selectable impact on organism fitness (Wingfield et al., 2006). Amplitudes are larger at higher latitudes (Fig 4D) and are likely to have stronger effects. From a clinical perspective, even a small systematic effect can cause misdiagnosis if the normal ranges are not adapted to the seasons, with associated costs of extra tests and treatment.

One test for the physiological relevance of the seasonal changes is to ask whether the coordinated effector hormone peak in winter/spring correlates with a time of high set-point for reproduction, metabolism, stress adaptation, and growth in humans. This relates favorably to observations on a winter-spring peak of human growth rate (Gelander, Karlberg & Albertsson, 1994; Land et al., 2005; Dalskov et al., 2016), cognitive functions (Meyer et al., 2016), immune functions (Dopico et al., 2015) and sperm quality (De Giorgi et al., 2015; Levitas 2013). Human fecundity also peaks in winter-spring in Israel and other countries in similar clines (Roenneberg and Aschof, 1990), and shifts to later in the year at higher latitudes. The relationship between hormone blood tests and fecundity is complex, since fecundity is affected by cultural factors. Nevertheless, the accumulated data suggests a winter/spring peak not only in effector hormones but also in the biological functions that they control.

The phases observed here, with spring delay in effector hormones and summer peaks of most pituitary hormones, are not accounted for by classic descriptions of the hormonal axes. These phases can be explained by the action of the hormones as growth factors for their downstream glands. This model may also explain seasonal patterns found in animal studies. Studies in diverse species find that effector hormones peak with a delay of few months following the shortest or longest photoperiod (Walkden-Brown et al., 1994; Walkden-Brown et al., 1997; Monfort et al., 1993). An approximate antiphase is found between effector and pituitary hormones in the HPA axis in several species including rams, horses, sand rats and beavers (Cordero et al., 2012; Ssewannyana et al., 1990; Czerwinska et al., 2015; Amirat and Brudieux, 1993). In agreement with the model predictions, the adrenal mass (Amirat et al., 1980) and anterior pituitary mass (Amirat and Brudieux, 1993) were found to change with seasons in a way that correlated with their hormonal output. Such as antiphase is not found, however, in studies on the male reproductive axis in goats (Walkden-Brown et al., 1994; Walkden-Brown et al., 1997). One marked difference between the present study and animal studies is that animals generally have larger seasonal amplitudes ranging from changes of tens to hundreds of percents.

Further tests of the present model can measure gland volumes with seasons, and more specifically the total functional masses of the hormone-secreting cell types. It is likely that gland-mass changes operate together with other slow processes such as epigenetic regulation to provide seasonal inertia. The gland-mass model is in line with wider evidence from animal studies, reviewed in (Piersma and Lindström, 1997), that indicate that the functional size of organs and aspects of the metabolic physiology of an individual may show flexibility over timescales of weeks and even days depending on physiological status, environmental conditions and behavioral goals.

This study used large-scale electronic medical records to obtain statistical power that allows a comprehensive view of hormone seasonality at the level of a few percent. To achieve this required filters to address ascertainment bias, including internal controls to remove records from people with illness or medications that affect each hormone, and to address circadian effects. Agreement with smaller-scale seasonality studies on cohorts of healthy individuals, where available, strengthens the confidence in this approach.

This study raises the possibility of an internal seasonal clock in humans that provides a high endocrine set-point for reproduction, growth, metabolism and stress adaptation in winter-spring. It suggests that large-scale medical records can be used to test mechanistic endocrine models and to provide a testing ground for dynamical physiological adaptations in humans.

## Supporting information

Supportive information

## Acknowledgements

Data were provided in part by the Human Connectome Project, WU-Minn Consortium (Principal Investigators: David Van Essen and Kamil Ugurbil; 1U54MH091657) funded by the 16 NIH Institutes and Centers that support the NIH Blueprint for Neuroscience Research; and by the McDonnell Center for Systems Neuroscience at Washington University.

We thank Benjamin Glaser, Shai Fuchs, Gil Levkowitz, Jacques Drouin, Patrice Mollard, Paul Le-Tissier, Johannes Dietrich, Ruslan Medzhitov and members of our labs for fruitful discussions, and Gabi Barabash and Ran Balicer for the Clalit-Weizmann collaboration. UA is the incumbent of the Abisch-Frenkel chair.

## Author contributions

Conceptualization, Av.T., A.B., N.M.C, O.K., Y.K., L.M., T.M., M.R., A.M., Am.T. and U.A.; Software, Av.T., A.B., N.M.C., Am.T., and U.A.; Methodology, Av.T. and U.A.; Formal Analysis, Av.T., A.B., N.M.C., Am.T., and U.A.; Investigation, Av.T., A.B., N.M.C., Am.T., and U.A.; Writing – Original Draft, Av.T. and U.A.; Writing – Review & Editing, Av.T., A.B., N.M.C, O.K., Y.K., L.M., T.M., M.R., A.M., Am.T. Supervision, U.A.; Funding Acquisition Am.T. and U.A.

## Methods

### Equations for the HPA model

The classic HPA hormone cascade with the feedback by cortisol on upstream hormones (Vinther *et al*., 2011) can be described by:

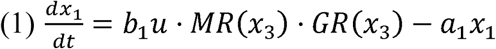

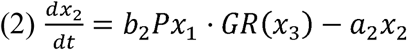

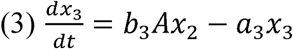

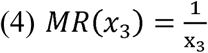

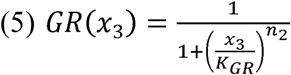

Where the concretion of CRH is *x*_1_, ACTH is *x*_2_ and cortisol is *x*_3_. *u* is the stress input to the hypothalamus (combined effect of psychological, circadian, seasonal and physiological stresses), *b*_*i*_ are the secretion rates, *a*_*i*_ the hormone removal rates, *P* and *A* are the ‘gland masses’ namely the functional mass of ACTH-secreting pituitary corticotrophs and cortisol-secreting adrenal cortex cells respectively. The equations model the negative feedback in which cortisol provides negative feedback on CRH secretion through MR, and negative feedback on both CRH and ACTH secretion through GR. Equation [4] describes the approximation that MR is at saturation ([*CORT*] ≫ *K*_*MR*_), as is commonly assumed under physiological conditions (Andersen,Vinther & Ottesen,2013; De Kloet *et al*. 1998).

In this study we use the approach of Karin et al (Karin et al., 2020) that adds two new equations to describe the effect of the hormones on the gland sizes. CRH activates P proliferation, and ACTH activates A proliferation:

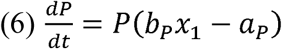

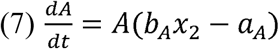

*a*_*P*_, *a*_*A*_ are the cell removal rates, and *b*_*P*_, *b*_*A*_ are the hormone-dependent growth rates. Equations for the timescale of months can be derived by a quasi-steady-state approximation for the hormones, showing a negative feedback loop between and *P* and *A*

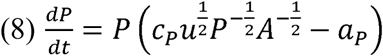

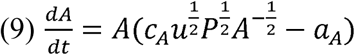

Where *c*_*P*_ and *c*_*A*_ are combinations of the parameters (SI section 6).

### Modelling

Input to most axes is thought to be dependent on photoperiod (Wehr et al., 1993) Photoperiod dependence on latitude, ***W***, was computed by the astronomical sunrise equation (SI section S7). The relative change in input to the model was proportional to the relative change in day length *u*(*t*) = 1 + *g* W(*ϕ*,)cos (*ωt*), where 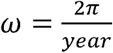, with maximal input at December 21 and amplitude W (day-length variation of 6-18 hours corresponds to W= 0.5). Unless noted otherwise, simulations where done to model Israel’s latitude *ϕ*, = 31.8° (day-length variation of approximately 10-14 hours, corresponds to W = 0.17). The factor g maps the photoperiod to the HP-axis input. We used g=0.5, which best fits the latitude dependence of cortisol measurements in Fig 4F. The value of g affects the amplitude but does not measurably affect the phases of the hormone dynamics (SI section S7). The amplitude of P with these parameters is 7.3% (Fig 4F). We simulated 4 years of seasonal input to avoid transients.

In order to simulate the response to a varying photoperiod, we simulated the fast equations (Eqs. 1-5) to obtain the numeric quasi-steady-state solution *x*_1,*qst*_, *x*_2,*qst*_, *x*_3,*qst*_ for a given input *u, P, A*. Next, we used these quasi steady-state solutions in Eq 6,7 to find the seasonally varying values of A and P. To simulate morning cortisol blood tests, we assumed based on data in SI2 that a person’s circadian phase and other acute HPA inputs at the time of the test are season-independent, and thus computed the steady-state fast equation response to an input u=1, using the seasonally varying value of A and P. Simulations were done using Python. Turnover rates for pituitary and adrenal (and other effector glands) were 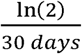, and for the liver 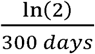 due to its longer turnover time of about a year (Nolan, Thomas and Levy, 2004; MacDonald, 1961).

### Data and code availability statement

The data and the code used in this study are available at https://github.com/alonbar110/Human-hormone-seasonality.

### Electronic Medical record data

The Clalit dataset contains the electronic health records (EHR) of 3.45 million individuals per year on average (Balicer and Afek, 2017). Data was anonymized by hashing of personal identifiers and addresses and randomization of dates by a random number of weeks uniformly sampled between 0 and 13 week for each patient and adding it to all dates in the patient diagnoses, laboratory and medication records. This approach maintained differential data analysis per patient. Diagnosis codes were acquired from both primary care and hospitalization records, and were mapped to the ICD9 coding system. The full study protocol was approved by the Clalit Helsinki Committee 0195-17-COM2.

### Data processing

We studied Clalit laboratory test records with more than total of 5000 tests (ACTH is the only test with a lower number, ∼4000 tests). For each test we analyzed data from all individuals with no chronic disease onset 6 months before the test, and no drug that affects the test bought in the 6 months before the test. Chronic disease was defined by non-pediatric ICD9 codes with a Kaplan-Meyer survival drop >10% over 5 years, and which are assigned above a minimal average rate of 1/3 per year. Drugs that affect a test were defined as drugs with significant effect on the test (FDR 0.01). Data was binned by months, correcting for date randomization by subtracting 6.5 weeks. The top and bottom 5% were removed rom each month bin to remove outliers. For each test, we then analyzed by quantiles per age decade bin for each gender. Seasonality was identified for each test by bootstrapping the data (with mean subtracted) and comparing to a zero-mean cosinor model with *A cos(ωt+*ϕ*) with ω=2π/year* against a null model of a constant level equal to zero. We also tested a second-order cosinor model *Acos*(*ωt* + *ϕ*) + *A*′ cos(2 *ωt* + *ϕ*′) for biannual effects, and selected the best model according to the Akaike information criterion. No tests justified a third-order or higher cosinor model. For each test, phase and amplitude were computed by bootstrapping (See SI). Processing of blood chemistry tests was identical to the hormone tests. Time of test for blood chemistry was also obtained in order to discern circadian patterns. Previous studies on cortisol (Fig 4D) were reanalyzed in terms of relative max-min amplitude. Error bars were computed by bootstrapping by months or semi-seasons. Details are in (SI section S11).

### Measurement of pituitary volume

Pituitary volumes were measured from MRI images from the Public Human Connectome Project (Glasser *et al*., 2013; Van Essen *et al*., 2013). We used 177 T1-weighted high-resolution 3T MR scans from the dataset “WU-Minn HCP”. Only this subset contained the image acquisition date and was useful for our seasonal analysis. To segment the pituitary we defined a region of interest (ROI) of 24×24×12 voxels at a fixed coordinates in all scans. The ROI was large enough to contain the pituitary and the hypothalamus in every subject. Manual pituitary segmentations used custom MATLAB software. Repeated segmentation of the same scan agreed well (intersection over union (IoU) mean = 0.895, std = 0.04). 5 scans were excluded from the analysis due to abnormal appearance. Volume of the pituitary (in cm^3^) was calculated by summing the areas of the voxels classified as pituitary tissue in the slices and multiplying by a factor of 1.6mm (slice thickness). Finally, pituitary volume was adjusted to represent its proportional volume to the intra-cranial volume (ICV). ICV values for each subject were obtained from the FreeSurfer (Fischl, 2012) datasheet supplied with the dataset. The manual pituitary segmentation was used to train an automated segmentation algorithm, based on the U-Net deep learning algorithm, a specialized convolutional neural network (CNN) architecture for biomedical image segmentation. Using a training set of 45 scans, the automated algorithm showed good agreement with manual segmentations on the remaining scans. Regression and 2-way ANOVA analysis indicates that seasonality in pituitary volume is not significantly affected by sex and age (SI section S12).

