## Supplementary material for "Human hormone seasonality": Supportive information

### Human hormone seasonality - Supporting Information

#### S1. Seasonality assessment of Clalit medical test database

We analyzed laboratory test data from the Clalit medical database, which included about 3.4 million active people followed from 2002 to 2017. For each test, we removed data from patients with chronic diseases and which bought drugs which affect the tests, as well as pregnant women. Clalit laboratory test records were studied in the age range of 20-80, with men and women analyzed separately. We show also results with age range 20-50 in Fig S1, which are generally very similar. For privacy reasons the date of each test was randomized by adding a number between 0 and 13 weeks, and we accordingly corrected the date by subtracting 6.5 weeks. For each test, we then transformed the raw measurement into quantiles per age decade bin for each gender. We then binned the tests into 12-month bins and computed the mean percentile value in each bin. Finally, we subtracted the overall mean percentile to result in percentile change as a function of month of the year.

To quantify seasonality of the percentile change, for each test $y$ we computed the oscillation amplitude and phase using a cosine fit, sometimes known as a zero-mean cosinor model (Russell *et al.*, 2008):

$$y\left( t \right)=Amp\cdot cos\left( \omega t+\phi\right)+e(t)$$

Where $Amp$ is the amplitude, $\phi$ is the phase and $e$ is the error of the data with respect to the cosinor model (Table S1). Error bars are computed by bootstrapping.

To identify seasonality, we compared the cosinor fit against a null model of a constant level equal to zero, y(t)=0. To test for biannual effects, we also tested a second-order cosinor model:

$$y\left( t \right)-M= Amp\cdot cos\left( \omega t+\phi\right)+{Amp}^{'}\cdot\cos\left( 2 \omega t+\phi^{'} \right) +e(t)$$

The best model was selected according to the Akaike information criterion. No tests justified a third-order or fourth-order cosinor model (not shown). For each test that identified seasonality, peak phase (acrophase) and relative max-min were computed by bootstrapping the error bar of each month.

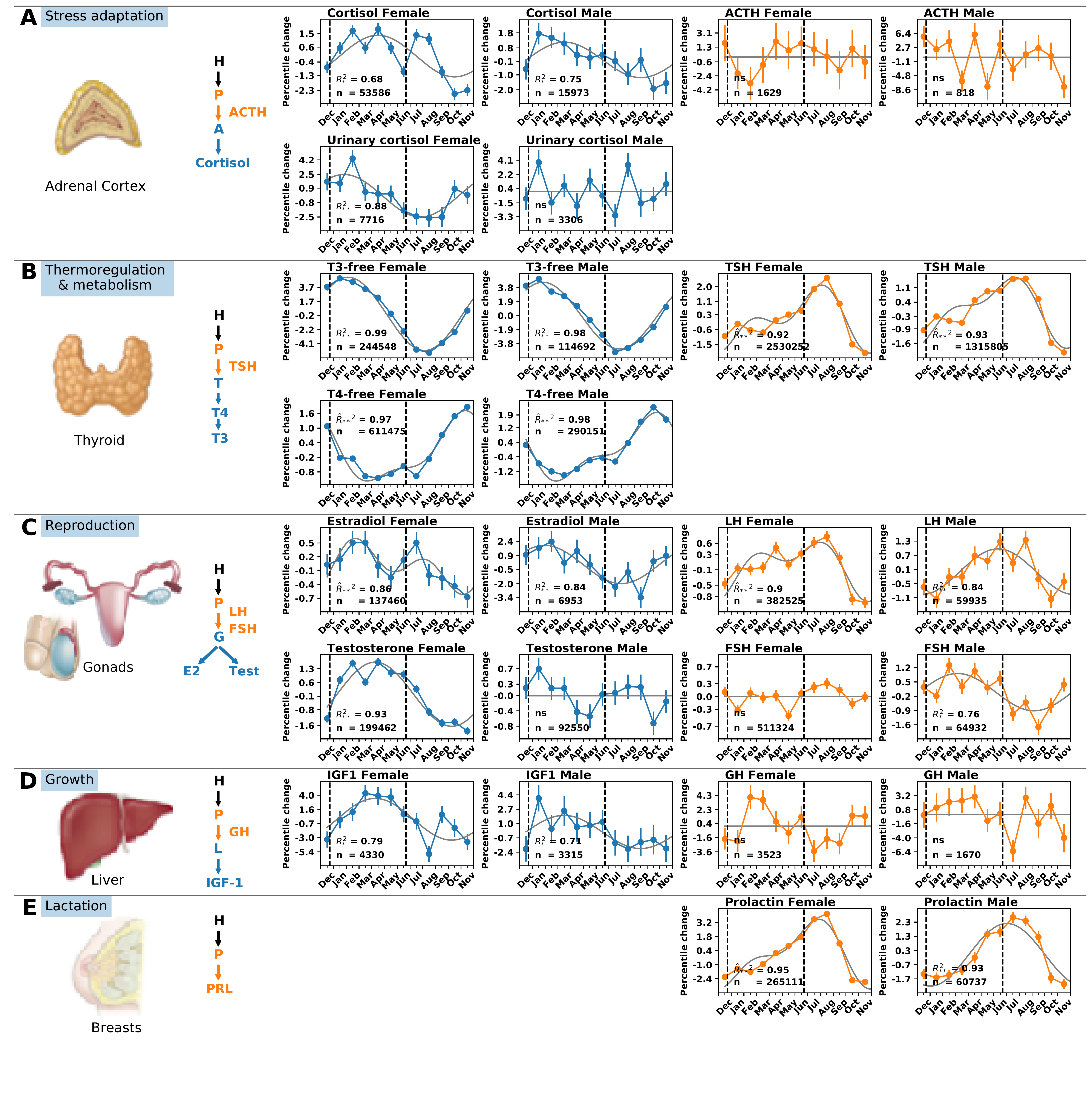

**Figure S1. Seasonality of hypothalamic-pituitary axes hormones from Clalit medical records.** Percentile change as a function of month of the year is shown, for data from the age range 20-50, for (A) HPA axis (B) Thyroid axis (C) Sex axis , (D) Growth axis (E) lactation pathway. Each panel indicates the number of tests n, zero-mean cosinor model (gray line) and $R^{2}$where significant ($R_{*}^{2}$ - $p<{5\cdot10}^{-2}$, $R_{**}^{2}$ - $p<{10}^{-3}$, ns - not significant), with first or second order model selected by Akaike criterion (second-order model is indicated by ^ above$R$). Vertical dashed lines indicate solstices Dec 21 and Jun21.

#### S2. Controls for circadian rhythm

Most hormones are secreted with a circadian rhythm (Kriegsfeld *et al.,* 2002). Thus, it is important to understand what time of day the tests were taken, especially for hormones with a short half-life. Among the hormone blood tests considered most sensitive to the time of day are tests for TSH, PRL, testosterone and cortisol. In contrast, the thyroid hormones T3 and T4 are considered insensitive due to the hormone lifetimes of one day and one week respectively. Tests that integrate over 24h such as urinary cortisol are considered less prone to circadian effects.

A major concern for the present study is whether the seasons affect the clock time of the tests. To address this, we obtained the clock time of each test in the Clalit database. We provide the distribution of the test times binned by four seasons for all tests and for males and females in Fig S2. The distributions are very similar between seasons, with a peak at 8am. We compared the distributions between each pair of seasons using a two-sample Kolmogorov–Smirnov test, corrected for multiple comparisons using FDR. We note that for some hormones, due to very high number of data-points, the K-S test reached significance although the effect sizes are minuscule (e.g. effect size 10-4 for TSH). We could not find any significant difference in test time distribution with effect size >0.01 (Fig S3). We therefore conclude that the distribution of test times in the Clalit database does not vary materially between the seasons.

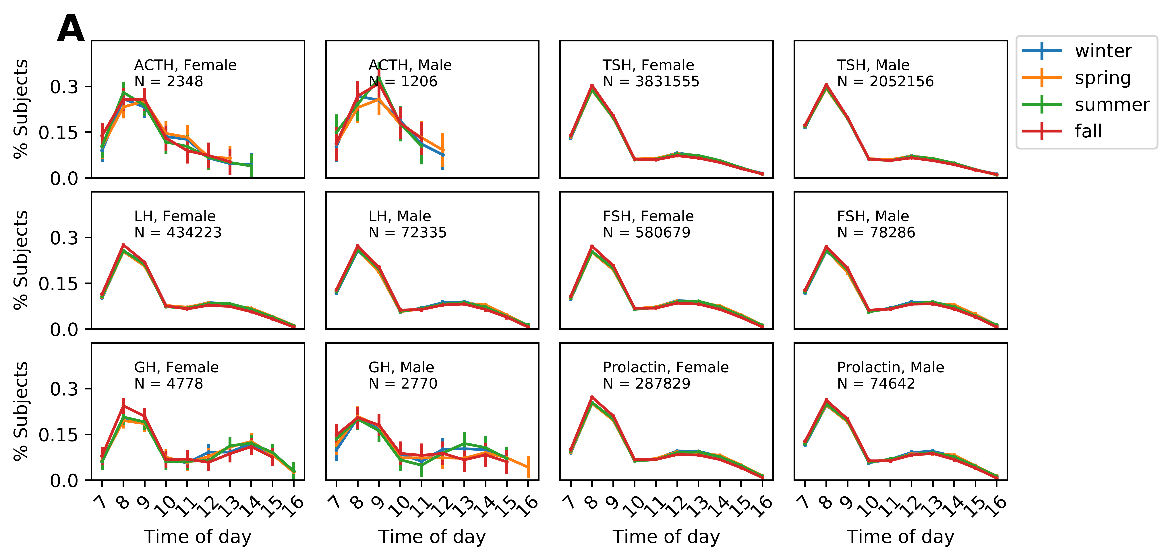

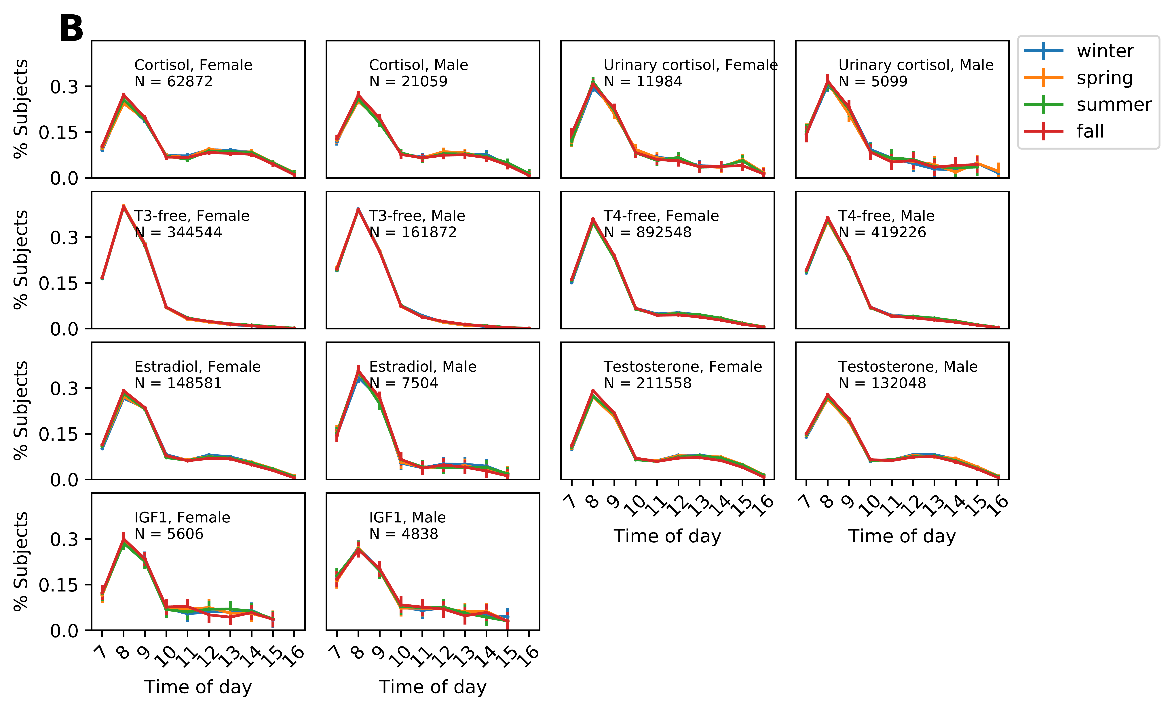

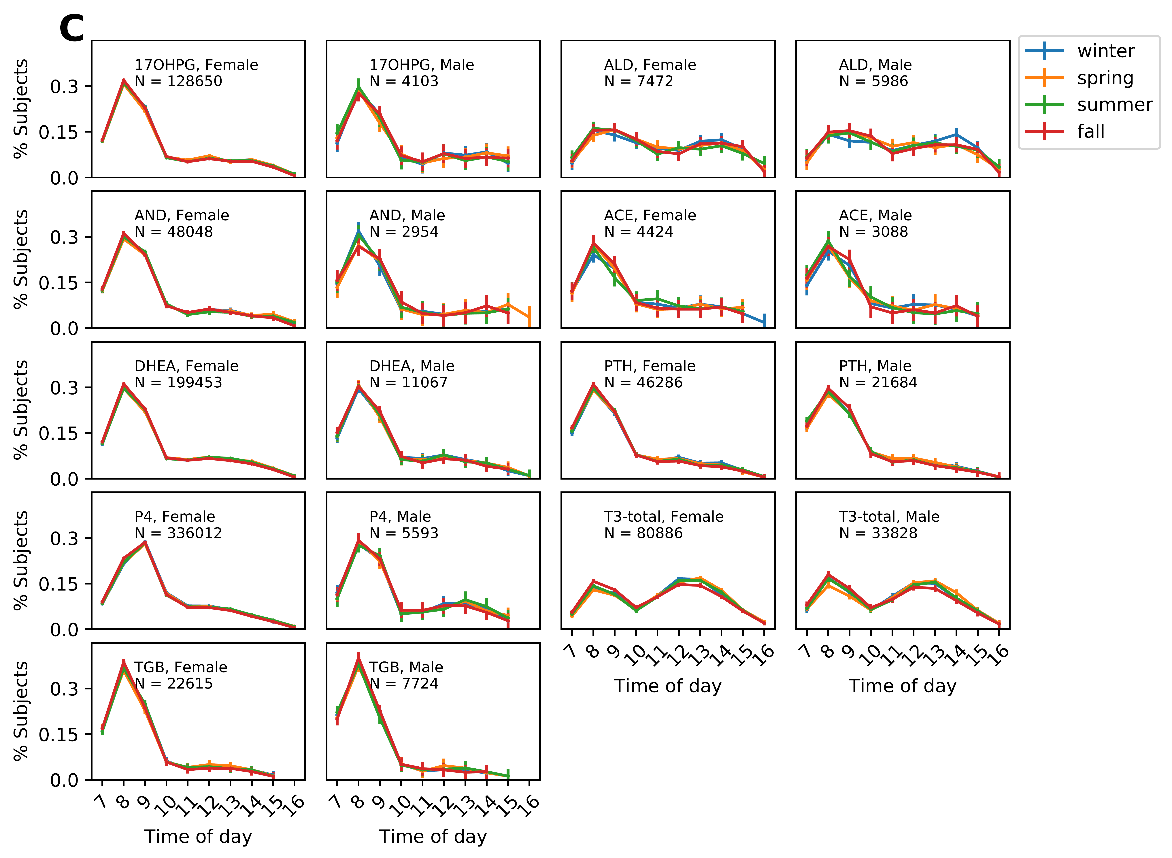

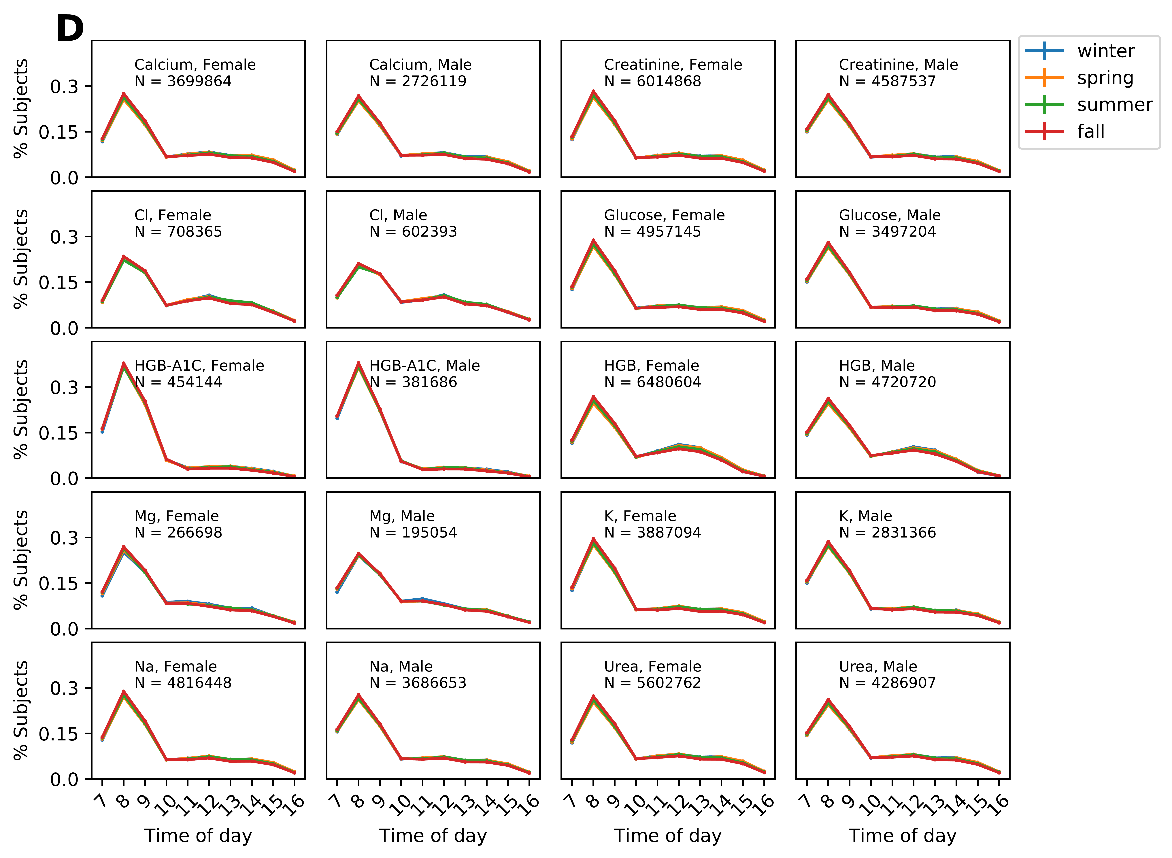

**Figure S2. Comparison of test time distribution in different seasons.** (A) Pituitary hormones. (B) Effector hormones. (C) Additional hormones that aren’t included in the main text. (D) Blood chemistry.

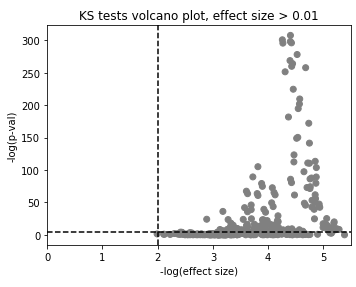
**Figure S3. Day-time distribution of tests done at different seasons are similar.** For each test, day-time distribution of every pair of seasons are compared. No comparison reach significance with effect size>0.01 (two-sample KS test)

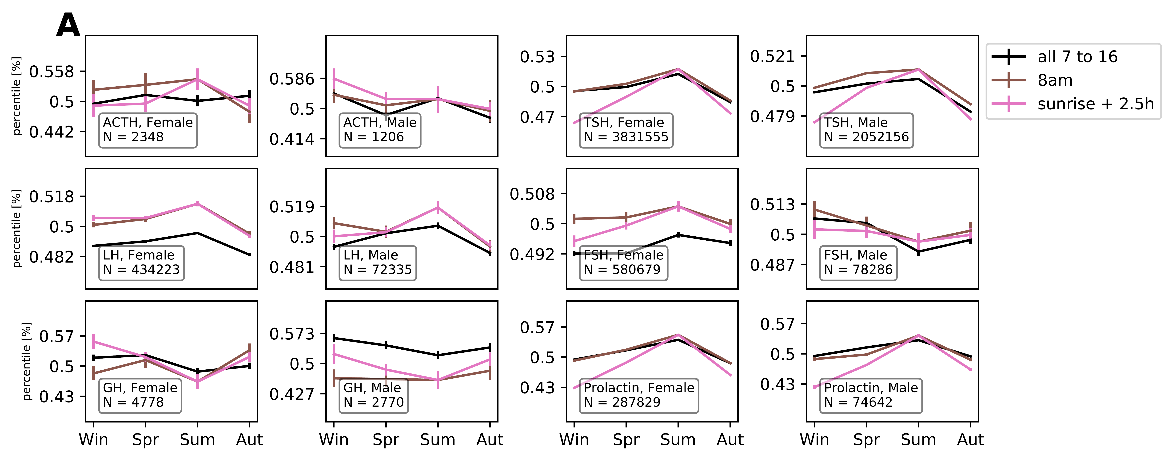

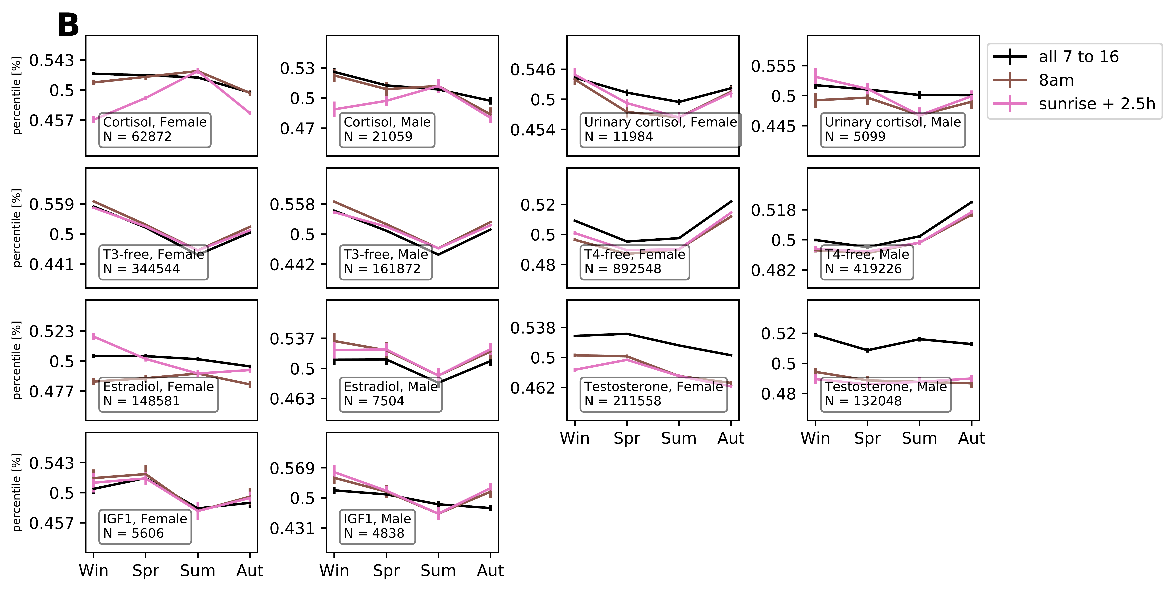

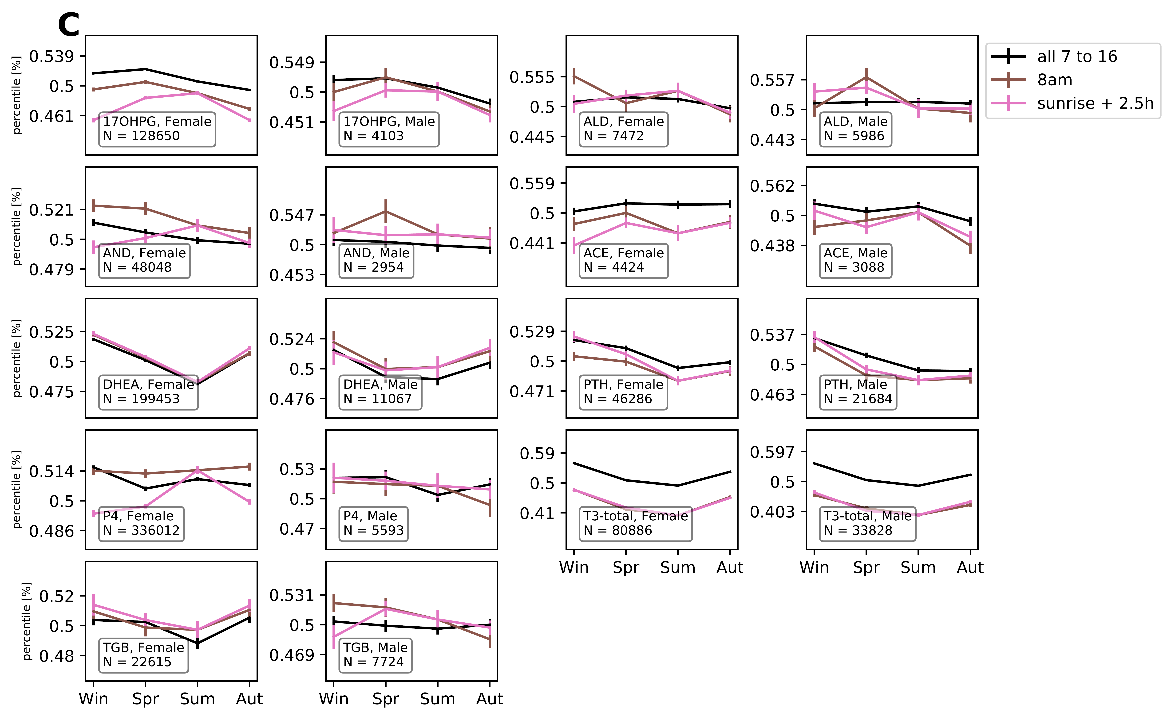

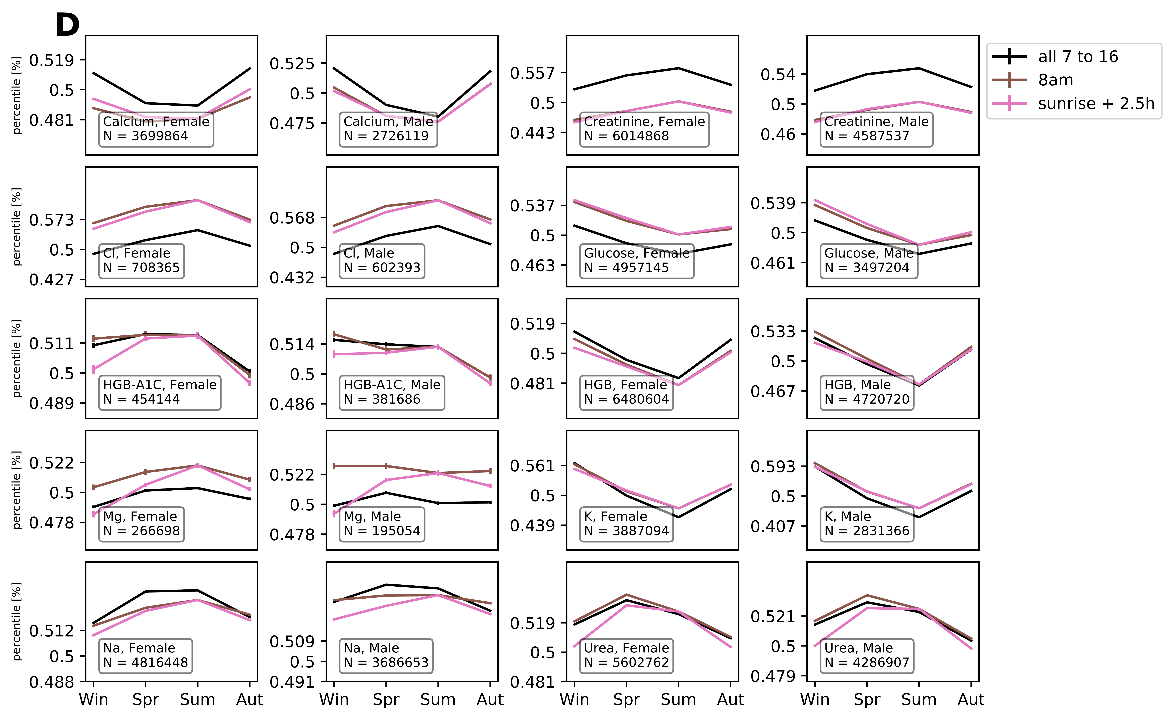

**Figure S4. Comparison of filters for hormone data.** (A) Pituitary hormones. (B) Effector hormones. (C) Additional hormones that aren’t included in the main text. (D) Blood chemistry.

We next used the clock time of each test to compare three filters. We compared mean percentile in four seasons of (i) tests done at all clock times, (ii) tests done between 8-9am, and (iii) tests done 2.5h(+/-0.5 h) after dawn, where time of dawn was computed for each season. The last filter aims to address concerns that tests done at a given time of day reflect different circadian phases relative to waking in different seasons. Most hormones with significant seasonality in Fig 1 (15/22) showed similar seasonal patterns with the same acrophase (correlation>0.7) in all three filters. Examples for T3 and for the three circadian-sensitive hormones TSH, PRL and Testosterone are shown in Fig S5. The exceptions (7/22 significant hormones) showed seasonality in filter iii that differed from filters i and ii, and are: cortisol blood m/f, testosterone m, estradiol f, GH m/f, ACTH f.

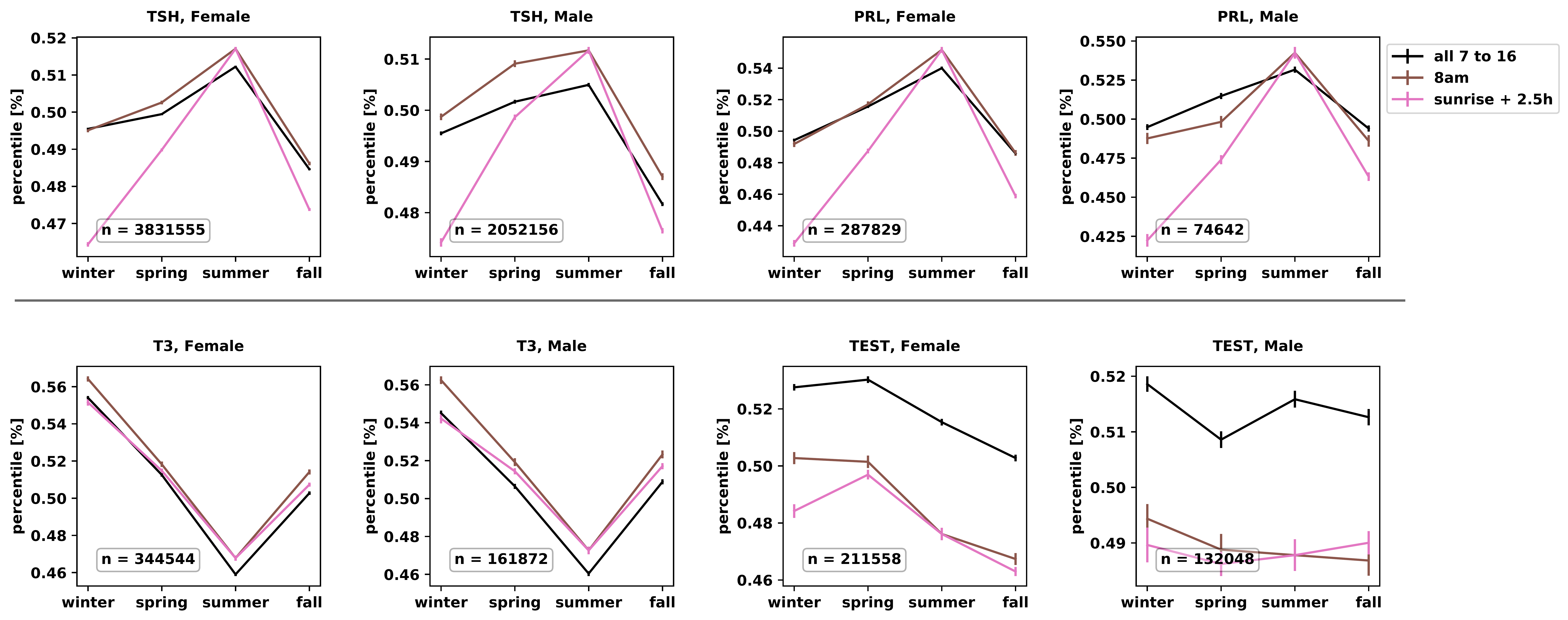

**Figure S5. Different methods to take time of day of tests into account generally agree.** For each test, three filters are compared: tests at all time of day (black line), tests only done between 8 and 9 am (brown line), and tests only done between 2 and 3 hours after dawn (pink line). Data is binned to four seasons. Each panel indicates hormone and male/female, as well as number of tests n.

One important test that showed sizable differences in seasonal behavior between the filters was blood cortisol. The results for filters i and ii (all times and 8 am) show a winter peak, whereas results for the dawn+2.5 hours filter (filter iii) do not. Since 24h urinary cortisol tests show a winter peak in all filters (Fig S6), and because most previous studies showed a winter peak for blood (Hadlow et al. 2018), hair (Abbel et al., 2016) and saliva (Persson et al., 2008) cortisol (where hair cortisol averages cortisol over at least one month and is expected to average over circadian rhythm well), we conclude that a consistent measurement for blood cortisol is the ‘all tests’ filter (filter i). We use this filter for all hormones in the main text.

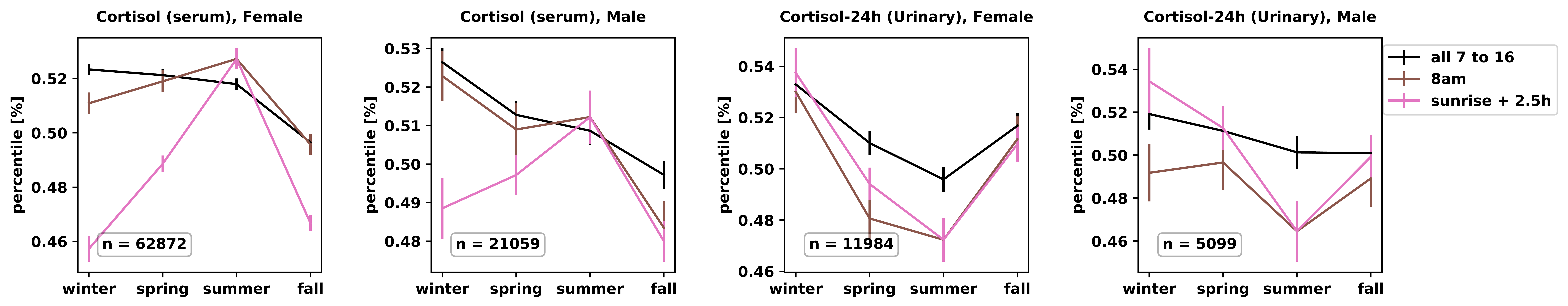

**Figure S6. Effect of time of day on cortisol tests.** For each test, three filters are compared: tests at all time of day (black line), tests only done between 8 and 9 am (brown line), and tests only done between 2 and 3 hours after dawn (pink line). Data is binned to four seasons. Each panel indicates the test (blood versus 24-h urinary cortisol) and male/female, as well as number of tests n.

#### S3. Seasonality of additional hormones in the Clalit medical database

We analyzed (Fig S7) additional hormones beyond those shown in Figure 1. This includes the reproductive-axis hormone progesterone (P4), androstenedione (AND) and 17-hydorxy-progesterone (17OHPG), as well as the adrenal hormones DHEA and aldosterone (ALD). We also analyzed several non-hypothalamic-pituitary blood tests such as parathyroid hormone (PTH) and angiotensin converting enzyme (ACE). Some of these do not show significant seasonality, perhaps due to the small sample size. Progesterone in females shows biannual seasonality similar to estrogen. PTH, AND and 17OHPG showed a spring-delay similar to other effector hormones.

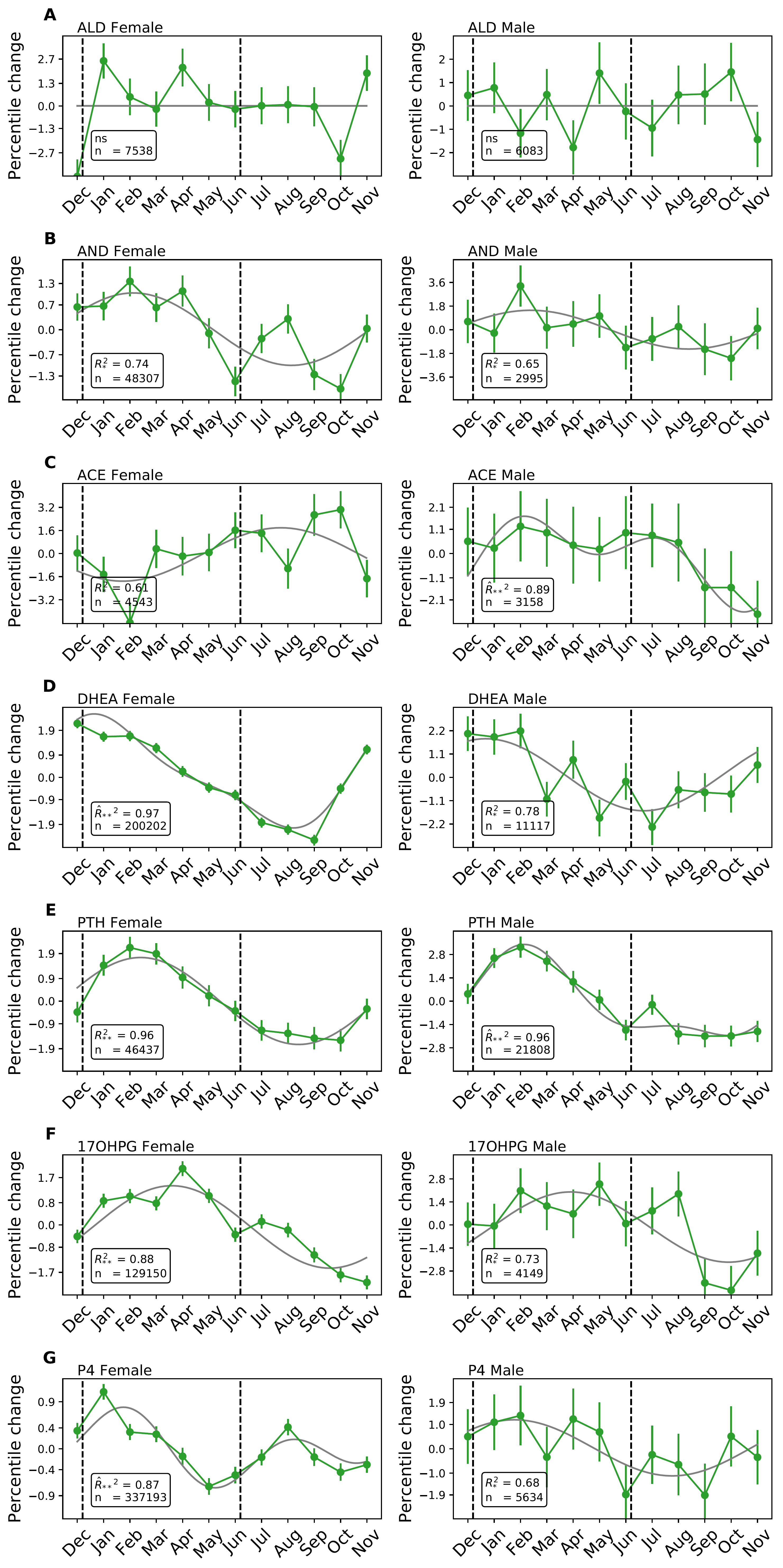

**Figure S7. Seasonality in additional hormones.** Green lines are the seasonal patterns of the hormones, gray lines are cosinor model fits. **(A)** Aldosterone. **(B)** Androstenedione. **(C)** Angiotensin-1 converting enzyme. **(D)** DHEA (Dehydroepiandrosterone). **(E)** Parathyroid hormone. **(F)** 17-hydroxy progesterone. **(G)** Progesterone. In each panel number of tests n, and cosinor model fit R^2 are indicated, with first or second order model selected by Akaike criterion. (Second order model is indicated by ^)

**Figure S8. Seasonality in blood chemistry tests.** Red lines are the data, gray lines are cosinor model fits. **(A)** Calcium. **(B)** Creatinine. **(C)** Chlorine. **(D)** Glucose. **(E)** Hemoglobin A1C. **(F)** Hemoglobin. **(G)** Magnesium. **(H)** Potassium. **(I)** Sodium. **(J)** Urea. In each panel number of tests n, and cosinor model fit R^2^ are indicated, with first or second order model selected by Akaike criterion. (Second order model is indicated by ^). Error bars are smaller than the marker size.

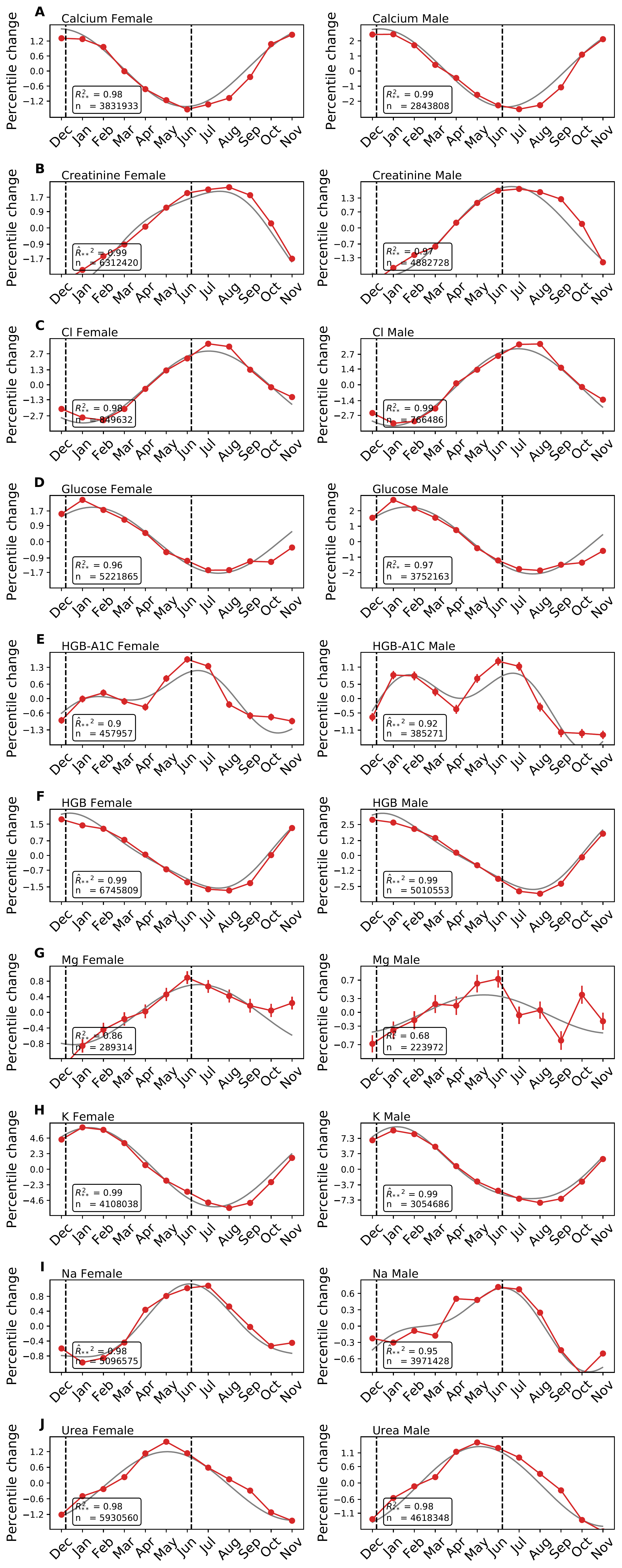

#### S4. All tests seasonality table

| Hormone | n | Sex | $1_{\mathrm{st}}\mathrm{Order}$  $phase \phi$  $[month]$ | $1_{\mathrm{st}}\mathrm{Order}$  $Amplitude Amp$  $[percentile]$ | $\mathrm{Acrophase}$  $[month]$ | Relative max-min amplitude [%] | $2_{\mathrm{nd}}\mathrm{Order}$  $\mathrm{phase}\phi^{'}$  $[month]$ | $2_{nd}\mathrm{Order}$  $Amplitude Amp$  $[percentile]$ |
| --- | --- | --- | --- | --- | --- | --- | --- | --- |
| ACTH | 2449 | female | 5.35±1.02 | 1.83±0.0074 | 5.58±1.52 | 4.15±0.0086 |  |  |
| ACTH | 1361 | male |  |  |  |  |  |  |
| TSH | 3920044 | female | 6.25±0.037 | 1.03±0.00018 | 8.0±4e-08 | 1.85±0.00033 | 3.79±0.042 | 0.86±0.00019 |
| TSH | 2095724 | male | 5.33±0.048 | 1.01±0.00026 | 7.07±0.26 | 1.45±0.00042 | 3.4±0.077 | 0.63±0.00027 |
| LH | 436231 | female | 5.93±0.21 | 0.54±0.00056 | 7.84±0.4 | 0.88±0.00085 | 4.17±0.31 | 0.37±0.00054 |
| LH | 72738 | male | 6.19±0.3 | 0.87±0.0014 | 7.56±0.86 | 1.35±0.002 |  |  |
| FSH | 583366 | female | 8.3±0.46 | 0.21±0.00047 | 7.78±0.82 | 0.46±0.00065 |  |  |
| FSH | 78711 | male | 2.69±0.29 | 0.92±0.0014 | 2.12±0.73 | 1.68±0.0024 |  |  |
| GH | 4821 | female | 2.85±0.56 | 2.01±0.0053 | 2.92±0.57 | 4.32±0.0071 |  |  |
| GH | 2786 | male |  |  |  |  |  |  |
| Prolactin | 289112 | female | 6.44±0.049 | 2.66±0.00068 | 7.99±0.076 | 3.31±0.0011 | 3.69±0.11 | 1.23±0.0007 |
| Prolactin | 75031 | male | 6.59±0.12 | 2.15±0.0013 | 7.42±0.5 | 2.34±0.0018 |  |  |
| Cortisol | 63175 | female | 3.74±0.23 | 1.3±0.0015 | 3.47±1.14 | 2.1±0.0022 |  |  |
| Cortisol | 21155 | male | 3.06±0.47 | 1.12±0.0027 | 1.54±1.02 | 2.11±0.0035 |  |  |
| Urinary cortisol | 12003 | female | 1.59±0.31 | 2.12±0.0035 | 1.88±0.5 | 3.07±0.0051 |  |  |
| Urinary cortisol | 5125 | male |  |  |  |  |  |  |
| T3-free | 345945 | female | 1.65±0.023 | 5.24±0.00063 | 1.01±0.076 | 5.32±0.0011 |  |  |
| T3-free | 162602 | male | 1.37±0.038 | 4.64±0.00087 | 1.0±-0.0 | 5.07±0.0015 |  |  |
| T4-free | 899982 | female | -0.73±0.048 | 1.57±0.00038 | 11.0±4e-08 | 1.76±0.0006 | 9.99±0.2 | 0.37±0.00039 |
| T4-free | 422709 | male | 10.15±0.069 | 1.51±0.0006 | 10.18±0.38 | 1.71±0.00087 | 9.58±0.18 | 0.61±0.00054 |
| Estradiol | 149177 | female | 3.46±0.5 | 0.38±0.00098 | 2.27±0.64 | 0.74±0.0013 | 3.6±0.6 | 0.34±0.00095 |
| Estradiol | 7554 | male | 1.66±0.44 | 1.9±0.004 | 2.3±1.84 | 3.14±0.0057 |  |  |
| Testosterone | 212613 | female | 3.74±0.095 | 1.6±0.0008 | 3.14±1.02 | 1.78±0.0012 |  |  |
| Testosterone | 132866 | male | -0.92±0.86 | 0.27±0.00092 | 1.06±0.27 | 0.88±0.0014 | 3.63±0.36 | 0.55±0.00096 |
| IGF1 | 5663 | female | 3.3±0.34 | 2.82±0.0052 | 3.59±0.99 | 4.85±0.0072 |  |  |
| IGF1 | 4891 | male |  |  |  |  |  |  |
| 17OHPG | 129150 | female | 3.58±0.13 | 1.47±0.001 | 4.0±0.096 | 2.06±0.0017 |  |  |
| 17OHPG | 4149 | male | 3.91±0.57 | 2.11±0.0056 | 4.67±2.47 | 4.03±0.0075 |  |  |
| ALD | 7538 | female |  |  |  |  |  |  |
| ALD | 6083 | male |  |  |  |  |  |  |
| AND | 48307 | female | 2.09±0.32 | 1.01±0.0017 | 2.49±1.16 | 1.68±0.0023 |  |  |
| AND | 2995 | male | 2.33±1.21 | 1.45±0.0063 | 2.14±1.6 | 3.62±0.0081 |  |  |
| ACE | 4543 | female | 7.74±0.62 | 1.86±0.0052 | 9.15±1.44 | 4.43±0.0083 |  |  |
| ACE | 3158 | male | 3.89±1.23 | 1.4±0.0061 | 3.94±2.96 | 3.29±0.0071 | 3.26±1.5 | 1.19±0.0054 |
| DHEA | 200202 | female | 1.3±0.077 | 2.05±0.00075 | 0.073±0.36 | 2.37±0.0013 | 0.11±0.27 | 0.6±0.00079 |
| DHEA | 11117 | male | 0.7±0.43 | 1.68±0.0035 | 1.03±1.08 | 2.78±0.0043 |  |  |
| PTH | 46437 | female | 2.41±0.19 | 1.72±0.0017 | 2.28±0.57 | 2.05±0.0024 |  |  |
| PTH | 21808 | male | 2.31±0.21 | 2.46±0.0025 | 1.91±0.55 | 3.04±0.0033 | 4.12±0.51 | 0.98±0.0024 |
| P4 | 337193 | female | 0.72±0.32 | 0.4±0.00059 | 1.0±-0.0 | 0.92±0.00096 | 3.91±0.24 | 0.48±0.00061 |
| P4 | 5634 | male | 1.66±1.03 | 1.17±0.0046 | 1.99±2.32 | 2.7±0.0057 |  |  |
| T3-total | 82998 | female | 0.82±0.066 | 4.03±0.0013 | 1.0±0.23 | 4.06±0.002 |  |  |
| T3-total | 34588 | male | 1.11±0.094 | 4.22±0.002 | 1.82±0.63 | 4.32±0.003 |  |  |
| TGB | 22733 | female | 1.21±0.51 | 0.98±0.0025 | 11.41±0.98 | 1.96±0.0033 | -0.25±0.6 | 0.86±0.0024 |
| TGB | 7820 | male |  |  |  |  |  |  |
| Ca | 3831933 | female | -0.088±0.022 | 1.55±0.00019 | 11.03±0.21 | 1.49±0.00032 |  |  |
| Ca | 2843808 | male | 0.35±0.017 | 2.58±0.00021 | 0.6±0.49 | 2.49±0.00034 |  |  |
| Creatinine | 6312420 | female | 6.76±0.012 | 2.47±0.00015 | 7.99±0.098 | 2.65±0.00026 | 6.22±0.056 | 0.5±0.00014 |
| Creatinine | 4882728 | male | 6.64±0.017 | 1.94±0.00016 | 6.92±0.27 | 2.05±0.00027 |  |  |
| Cl | 849632 | female | 7.04±0.027 | 3.12±0.00044 | 7.03±0.18 | 3.33±0.00069 |  |  |
| Cl | 766486 | male | 6.95±0.024 | 3.41±0.00046 | 7.63±0.49 | 3.54±0.00066 |  |  |
| Glucose | 5221865 | female | 1.49±0.018 | 1.83±0.00016 | 1.0±-0.0 | 1.96±0.00026 |  |  |
| Glucose | 3752163 | male | 1.66±0.018 | 2.16±0.0002 | 1.0±-0.0 | 2.28±0.00033 |  |  |
| HbA1c | 457957 | female | 5.26±0.12 | 0.91±0.00051 | 6.06±0.23 | 1.32±0.00075 | 1.94±0.18 | 0.59±0.00056 |
| HbA1c | 385271 | male | 4.29±0.12 | 0.91±0.00063 | 6.18±0.51 | 1.34±0.00089 | 2.45±0.14 | 0.82±0.00059 |
| HGB | 6745809 | female | 0.84±0.016 | 1.7±0.00014 | 0.0±-0.0 | 1.7±0.00023 | -0.62±0.097 | 0.3±0.00015 |
| HGB | 5010553 | male | 1.0±0.011 | 2.9±0.00016 | 0.0±-0.0 | 2.98±0.00028 | -0.55±0.061 | 0.55±0.00017 |
| Mg | 289314 | female | 6.56±0.18 | 0.77±0.00063 | 6.13±0.45 | 1.18±0.0011 |  |  |
| Mg | 223972 | male | 5.32±0.37 | 0.42±0.00075 | 5.69±0.81 | 0.79±0.0011 |  |  |
| K | 4108038 | female | 1.33±0.006 | 5.85±0.00019 | 1.0±-0.0 | 5.96±0.00033 |  |  |
| K | 3054686 | male | 1.27±0.0049 | 8.48±0.00022 | 1.0±-0.0 | 8.56±0.00037 | 2.02±0.033 | 1.16±0.00021 |
| Na | 5096575 | female | 6.22±0.032 | 0.96±0.00017 | 6.87±0.34 | 1.03±0.00027 | 0.055±0.13 | 0.23±0.00017 |
| Na | 3971428 | male | 5.2±0.057 | 0.62±0.00018 | 6.21±0.41 | 0.81±0.0003 | 1.65±0.13 | 0.27±0.00018 |
| Urea | 5930560 | female | 5.04±0.023 | 1.31±0.00015 | 5.0±-0.0 | 1.51±0.00025 |  |  |
| Urea | 4618348 | male | 5.12±0.023 | 1.46±0.00017 | 5.0±-0.0 | 1.62±0.0003 |  |  |

**Table S1.** Seasonality pattern of laboratory tests for various hormones. Error bars are form bootstrapping.

#### S5.Blood chemistry tests show seasonality with phases concentrated around the solstices

We analyzed the seasonality of blood chemistry tests. Fig S8 shows the percentile changes as a function of month of the year for males and females. We find seasonality for almost all tests, with very small error bars due to the large number of tests. Biannual seasonality is found for hemoglobin A1C (HGBA1C).

Blood chemistry tests tended to peak near the solstices Dec21 and Jun 21 (Fig 3). To quantify the tendency to peak near solstices, we computed for each test the absolute time difference $d$(modulo 12 months) between its peak (acrophase) time, $t_{peak}$ , and the closest solstice: $d=\min\left( \left| t_{peak}-Dec21 \right|,\left| t_{peak}-Jun21 \right| \right)$. The blood chemistry tests average about $d =0.75\pm0.11$months (mean $\pm$ SE), in contrast to the pituitary/effector hormones which average $d= 1.5\pm0.14$month. The distributions of *d* differ significantly between chemistry tests and pituitary/effector tests (effect size=1.58, KS test p<10^-10^).

The peak phases near solstices may suggest a simple model for many of these tests in which current seasonal input affects the blood chemistry variables, without delays caused by mechanisms such as the functional mass changes suggested for pituitary/effector hormones. Exceptions with a sizable delay include urea and creatinine, related in part to kidney functions. Similarly, glucose may show delays related to turnover of beta cells (Karin, 2016).

#### S6. Mass-change model parameters

We use the model of Karin et al (2020) as a minimal model of the HPA axis with changes in functional mass of the hormone-secreting cells. The model is based on the HPA model of Ottesen et al (2011) which was designed to describe the fast timescale of hours-days, and thus assumed that gland mass is constant. Karin et al added two equations to describe the changes in functional mass, by taking into account the effect of CRH as the main growth factor for corticotrophs, and ACTH as the main growth factor for adrenal cortex cells. Let $x_{1}$, $x_{2}$ and $x_{3}$ be CRH, ACTH and cortisol concentrations, respectively. P is the total functional mass of corticotroph cells that secrete ACTH and A is the total functional mass of adrenal cortex cells that secrete cortisol. The model equations are:

(1) $\frac{dx_{1}}{dt}=b_{1}u\cdot MR\left( x_{3} \right)\cdot GR\left( x_{3} \right)-a_{1}x_{1}$
(2) $\frac{dx_{2}}{dt}=b_{2}Px_{1}\cdot GR\left( x_{3} \right)-a_{2}x_{2}$
(3) $\frac{dx_{3}}{dt}=b_{3}Ax_{2}-a_{3}x_{3}$
(4) $\frac{dP}{dt}=P\left( b_{P}x_{1}-a_{P} \right)$
(5) $\frac{dA}{dt}=A(b_{A}x_{2}-a_{A})$
(6) $MR\left( x_{3} \right)=\frac{1}{x_{3}}$
(7) $GR\left( x_{3} \right)=\frac{1}{1+\left( \frac{x_{3}}{K_{GR}} \right)^{n}}$

The functions for GR and MR (Eq. 6-7) describe the inhibition of hormone secretion by cortisol through the glucocorticoid receptor (GR) and the mineralocorticoid receptor (MR). The MR receptor activity is modeled as 1/$x_{3}$ which approximate a Michaelis-Menten term 1/(𝑘+𝑥3) in the limit where hormones exceed their Michaelis constant 𝑥3≫𝑘. The low affinity and cooperative GR is modeled by a Hill function, with half-way effect at $K_{GR}$. The values of the parameters are provided in table S2. As stated in the main text, the $a_{i}$ parameters are the removal rates from the literature. We note that the present study focuses on the slow timescale of weeks, and the rates $a_{1},a_{2},a_{3}$ do not affect the results on this timescale. Here $a_{P},a_{A}$ are the cell removal rates, and $b_{P},b_{A}$ are the hormone-dependent proliferation rates.

| Parameter | Value | Reference |
| --- | --- | --- |
| $a_{1}$ | 0.17/min | (Andersen, Vinther and Ottesen, 2013) |
| $a_{2}$ | 0.035/min | (Andersen, Vinther and Ottesen, 2013) |
| $a_{3}$ | 0.0091/min | (Andersen, Vinther and Ottesen, 2013) |
| $a_{P}$ | log(2)/20day |  |
| $a_{A}$ | log(2)/30day |  |
| $K_{GR}$ | 4 |  |
| $n$ | 3 | (Andersen, Vinther and Ottesen, 2013) |
| $b_{1}$ | 0.17/min | Normalization of all variable steady-states to 1 |
| $b_{2}$ | 0.035/min | Normalization of all variable steady-states to 1 |
| $b_{3}$ | 0.0091/min | Normalization of all variable steady-states to 1 |
| $b_{P}$ | log(2)/30day | Normalization of all variable steady-states to 1 |
| $b_{A}$ | log(2)/30day | Normalization of all variable steady-states to 1 |

**Table S2. Parameter values.**

#### S7. Analysis of model properties

We used a simplified form of the model for analytical analysis. We use the fact that physiological cortisol levels are usually too low to appreciably activate GR, and retain only the effect of MR in the feedback loop (cortisol feeds back on hypothalamic CRH secretion through MR, whereas ACTH secretion is regulated by GR but not by MR):

(8) $\frac{dx_{1}}{dt}=\frac{b_{1}u}{x_{3}}-a_{1}x_{1}$
(9) $\frac{dx_{2}}{dt}=b_{2}Px_{1}-a_{2}x_{2}$
(10) $\frac{dx_{3}}{dt}=b_{3}Ax_{2}-a_{3}x_{3}$
(11) $\frac{dP}{dt}=P(b_{P}x_{1}-a_{P}\boldsymbol{)}$
(12) $\frac{dA}{dt}=A(b_{A}x_{2}-a_{A}\boldsymbol{)}$

##### Derivation of quasi-steady-state equations for the HPA model

The steady-state hormone levels (averaged over months) in this model is robust to almost all model parameters:

(13) $x_{1,st}=\frac{a_{P}}{b_{P}}$
(14) $x_{2,st}=\frac{a_{A}}{b_{A}}$
(15) $x_{3,st}=\frac{b_{1}b_{P}}{{a_{1}a}_{P}}u$

Robustness of hormone baseline levels originates from the integral-feedback terms in Eq. 4-5. In contrast, the classic model without gland mass changes (Eq 1-3,6-7 with constant A and P, and without Eq 4-5) has a steady-state that depends on more parameters such as hormone removal and production parameters:

(16) $x_{1,qst}=\left( \frac{b_{1}a_{2}a_{3}}{a_{1}b_{2}b_{3}}u^{1}P^{-1}A^{-1} \right)^{\frac{1}{2}}$
(17) $x_{2,qst}=\left( \frac{b_{1}b_{2}a_{3}}{a_{1}a_{2}b_{3}}uPA^{-1} \right)^{\frac{1}{2}}$
(18) $x_{3,qst}=\left( \frac{b_{1}b_{2}b_{3}}{a_{1}a_{2}a_{3}}uPA \right)^{\frac{1}{2}}$

These terms also serve as quasi-steady state solutions of the full model (eq 1-7) on the short timescale of hours. Equations for the full model on a timescale of months can be derived by using the quasi-steady-state approximation for the hormones, and substituting it into Eq 4-5. This results in two coupled ODEs for the functional masses A(t) and P(t):

(19) $\frac{dP}{dt}=P(c_{P}u^{\frac{1}{2}}P^{-\frac{1}{2}}A^{-\frac{1}{2}}-a_{P})$
(20) $\frac{dA}{dt}=A(c_{A}u^{\frac{1}{2}}P^{\frac{1}{2}}A^{-\frac{1}{2}}-a_{A})$

Where:

$c_{P}=\left( \frac{b_{1}a_{2}a_{3}}{a_{1}b_{2}b_{3}} \right)^{\frac{1}{2}}b_{P}$, $c_{A}=\left( \frac{b_{1}b_{2}a_{3}}{a_{1}a_{2}b_{3}} \right)^{\frac{1}{2}}b_{A}$

In the case where we normalize the steady-state of the variables to one, we have $b_{i}=a_{i}$, and hence $c_{P}=a_{P}$ and $c_{A}=a_{A}$. This system of equation describes a nonlinear negative feedback loop, in which $P$ increases $A$ and $A$ reduces $P$.

##### Linear stability analysis of the fixed point, and conditions for a spiral fixed point

This slow timescale equations are a special case of

(21) $\frac{dP}{dt}=P(c_{P}u^{n_{uP}}P^{n_{PP}}A^{n_{AP}}-a_{P})$
(22) $\frac{dA}{dt}=A(c_{A}u^{n_{uA}}P^{n_{PA}}A^{n_{AA}}-a_{A})$

Models of this form were analyzed by Komarova et al (2003) in the context of bone cell circuits. If the system is nondegenerate, it has a unique nonzero fixed point, which we denote $\left( P_{st}, A_{st} \right)$. To check the behavior around the fixed point, we compute the Jacobian (the matrix of derivatives at the fixed point):

(23) $\left( \begin{matrix} \frac{d\dot{P}}{dP} & \frac{d\dot{P}}{dA} \\ \frac{d\dot{A}}{dP} & \frac{d\dot{A}}{dA} \end{matrix} \right)=\left( \begin{matrix} n_{PP}a_{P} & n_{AP}a_{P}\frac{P_{st}}{A_{st}} \\ n_{PA}a_{A}\frac{A_{st}}{P_{st}} & n_{AA}a_{A} \end{matrix} \right)$

To determine the stability of the fixed point we compute the determinant and trace of the Jacobian:

(24) $det=a_{P}a_{A}(n_{PP}n_{AA}-n_{AP}n_{PA})$
(25) $trace={a_{P}n}_{PP}+a_{A}n_{AA}$

Here:$n_{PP}=-\frac{1}{2}, n_{AP}=-\frac{1}{2},n_{PA}=\frac{1}{2},n_{AA}=-\frac{1}{2}$. Thus: $det>0$and$trace<0$ (note that $a_{P},a_{A}>0$). Therefore, the fixed point is stable for any values of tissue turnover timescale$a_{P},a_{A}$. The fixed point is a spiral if ${trace}^{2}<4det$ . To determine the type of fixed point, we compute:

(26) $trace^{2}-4det= a_{P}^{2}(a_{R}^{2}n_{AA}^{2}+(4n_{AP}n_{PA}-2n_{PP}n_{AA})a_{R}+n_{PP}^{2})$

Here, $a_{R}=a_{A}/a_{P}$ is the ratio between the timescale of the glands. Substituting the values of the exponents yield:

(27) $trace^{2}-4det=a_{P}^{2}(\frac{1}{4}a_{R}^{2}-\frac{3}{2}a_{R}+\frac{1}{4})$

This expression is negative if: $3-\sqrt{8}<a_{R}<3+\sqrt{8}$, which is about a 34-fold range, $0.17<a_{R}<5.8$ . Thus, if the timescales of the two glands are similar enough (that is, if$a_{R}$is close to one), the fixed point is spiral and can support damped oscillations (Strogatz, 2015).

##### Noise driven sustained oscillations

Noise in the system can cause sustained oscillations (Geva-Zatorsky, et al., 2010; Alon, 2019) with frequency approximately equal to the resonance frequency of the spiral fixed point, $\boldsymbol{\omega}_{\mathbf{0}}=\sqrt{\boldsymbol{det}-\frac{\boldsymbol{trac}\boldsymbol{e}^{\mathbf{2}}}{\mathbf{4}}}$ (Fig S9). The noise represents stresses and physiological variations. Such sustained oscillations suggest a model for the intrinsic circa-annual clock revealed by experiments in which animals are kept under constant photoperiod and temperature conditions (Zucker, 2001; Lincoln et al., 2006; Gwinner, 2012). For orientation, note that when the timescales of the two tissues are equal, the resonance frequency is half of the tissue removal rate parameter$, \boldsymbol{\omega}_{\mathbf{0}}=\frac{\boldsymbol{a}_{\boldsymbol{P}}}{\mathbf{2}}$. The resonance period is equal to one year in this case when turnover time is on the order of one month: ${\boldsymbol{a}_{\boldsymbol{A}}=\boldsymbol{a}}_{\boldsymbol{P}}=\mathbf{4}\boldsymbol{\pi}\left[ \frac{\mathbf{1}}{\boldsymbol{year}} \right]=\frac{\mathbf{1}}{\mathbf{29}}[\frac{\mathbf{1}}{\boldsymbol{day}}]$. Cell half-lives in this case are $\frac{\log\left( \mathbf{2} \right)}{\boldsymbol{a}_{\boldsymbol{P}}}=\mathbf{20}$days. Such a timescale is roughly consistent with experiments on turnover of corticotrophs and adrenal cortex cells in rodents (Westlund, Aguilera and Childs, 1985; Gulyas, Puztai and Makara, 1991). ”

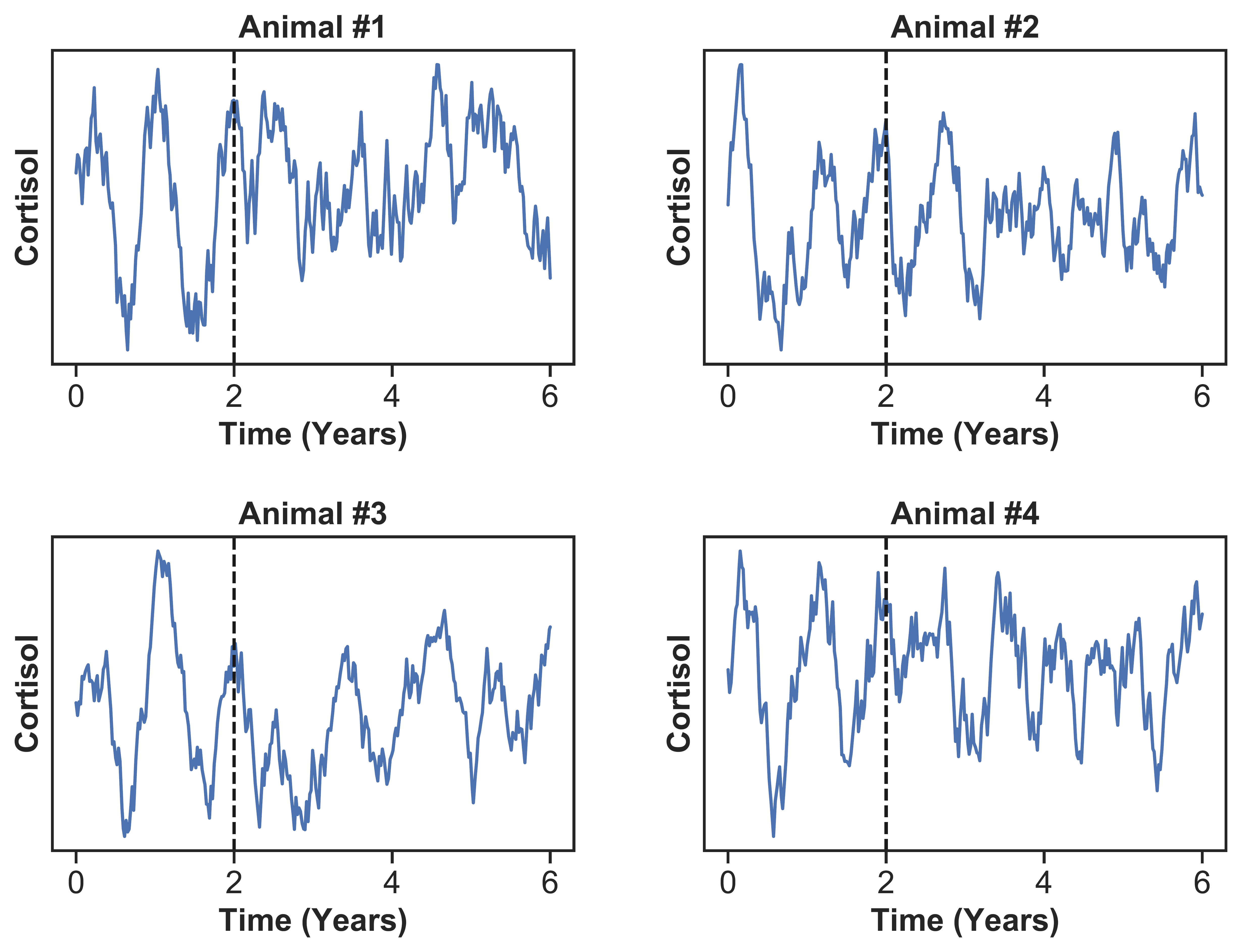

**Fig S9. HPA model with gland mass changes produces noise-driven sustained oscillations.** HPA model was simulated for 6 years. For the first 2 years, the input included seasonal (day length) component plus noise. In the next 4 simulated years, the seasonal component was replaced with constant value representing constant photoperiod conditions. The vertical dashed line indicate transition into constant photoperiod conditions. Each subplot is an example of one simulation. Input was $u\left( t \right)=1+g*u_{0}(\phi)cos\left( \omega t \right)+noise$ during the first two years and $u\left( t \right)=1+noise$ in the next 4 years, where $\phi={50}^{o}$, $g=0.5$, $\omega=\frac{2\pi}{year}$. Noise was piecewise constant: a uniformly distributed random number$U\left( \left[ 0.5,1.5 \right] \right),$ with a different number generated each simulated week.

##### Sensitivity of hormone phases to model parameters

The seasonal phases in the HPA model are nearly insensitive to changes in any of the hormone production and removal parameters. Doubling or halving each parameter didn’t measurably changes the phase of peak cortisol. The phases depend strongly only on the two tissue removal rate parameters, of corticotroph and adrenal cortex cells. These parameters are $a_{P}$ and $a_{A}$. The cell half-lives are $\frac{\log\left( 2 \right)}{a_{P}}$ and $\frac{\log\left( 2 \right)}{a_{A}}$. We analytically solved for the effect of changing these parameters on the phases of ACTH and cortisol (Fig S10). We find that ACTH phase (Apr-Aug) is captured by adrenal half-lives of under 30 days. The cortisol phase (Jan-Mar) is captured by a region in parameter space where both adrenal and corticotroph half-lives exceed 15 days. Both phases are captured simultaneously by a region in parameter space. This region also intersects the parameters that provide a resonance period of around 1 year (10-14 months) in the linearized model equations around the spiral fixed point. This indicates estimated half-lives of 25 days (turnover of 17 days) for cortisol secreting adrenal cells and 40 days (turnover of 27 days) for pituitary corticotrophs that secrete ACTH. We conclude that there is a sizable parameter range (dark green region in Fig S10) in which the phases match the observed Clalit tests data. Similar dependence on parameters is found for the other HP-axis models described in this SI.

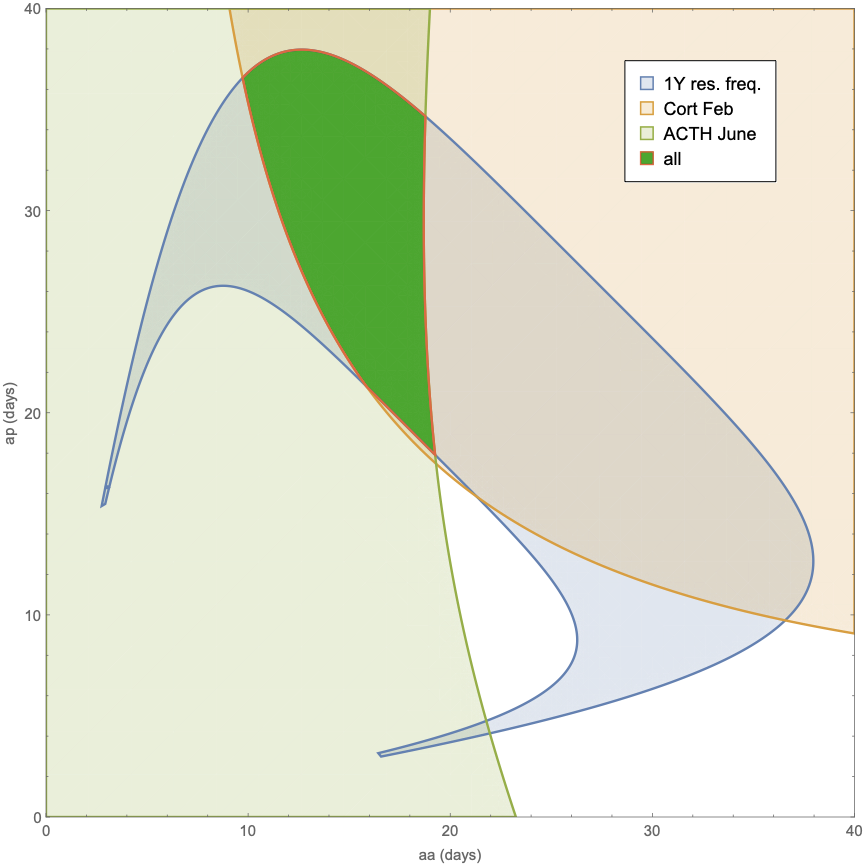

**Fig S10: The HPA hormone phases are captured by the HPA model with tissue turnover times in the range of weeks.** Analytical solution of the linearized model for the range in parameter space (turnover parameters for corticotrophs and adrenal cells) in which the experimentally observed phases are captured by the model. Light green region: ACTH acrophase April-August. Light red region: cortisol phase Jan-Mar. Light blue region: parameters that provide a resonance frequency to the spiral fixed point of about a year (10-14 months). These regions intersect in the dark green region.

#### S8. Photoperiod input to the HPA model

We model the seasonal input to the HPA axis according to the change in photoperiod across seasons. This change increases with latitude according to the sunrise equation (Teets, 2003):

(28) $\cos\left( \alpha_{0} \right)=-\tan\left( \phi\right)tan(\delta)$

Where $\alpha_{0}$ is the hour angle of sunrise (or sunset), $\phi$ is the latitude and $\delta=23.44^{\circ}$ is the earth’s tilt. From the sunrise equation we can derive the length of the longest day from sunrise to sunset:

(29) $\frac{2}{{15}^{\circ}}\cos^{-1} (\tan\left( \delta\right)\tan\left( \phi\right)) [hour]$

From this we compute the relative change in photoperiod from 12 hours at the longest or shortest day:

(30) $u_{0}\left( \phi\right)=\frac{12-\frac{2}{{15}^{\circ}}\cos^{-1} (\tan\left( \delta\right)\tan\left( \phi\right))}{12}$

We note that the amplitude of the photoperiod signal might be transformed by the brain (for example, by the melatonin system). To model this, we introduce a gain factor $g$ to obtain the effective input to the hypothalamus. The input thus varies with seasons as

(31) $u\left( t \right)=1+gu_{0}\left( \phi\right)cos(\omega t)$

with $\omega=2\pi/year$. We use $g$= 0.5 to obtain good agreement with cortisol data (Fig 4D). We note that latitude has a small effect on the hormone phases; cortisol delay from winter solstice is smaller at latitude ${60}^{o}$ by about 5% than at ${20}^{o}$.

#### S9. Models for additional HP-axes

We extended the minimal-model approach used for the HPA axis above to the other pituitary axes. We used experimentally described interactions in all models, including the effect of the hormones as growth factors for the tissues.

##### **HPG axis**

We model the HPG (hypothalamic-pituitary-gonadal) axis with the feedback by sex steroids on upstream hormones, together with gland mass changes, as follows:

(32) $\frac{dx_{1}}{dt}=\frac{b_{1}u}{x_{3}}-a_{1}x_{1}$
(33) $\frac{dx_{2}}{dt}=\frac{b_{2}P_{G}x_{1}}{x_{3}}-a_{2}x_{2}$
(34) $\frac{dx_{3}}{dt}=b_{3}Gx_{2}-a_{3}x_{3}$
(35) $\frac{dP_{G}}{dt}=P_{G}(b_{GT}x_{1}-a_{GT}\boldsymbol{)}$
(36) $\frac{dG}{dt}=G(b_{G}x_{2}-a_{G})$

Here hormone concentration of GnRH is $x_{1}$ (note that in this axis the pulse frequency of GnRH may be decisive and $x_{1}$ can be taken to describe this frequency), LH /FSH is $x_{2}$ and Testosterone/Estradiol (T/E) is $x_{3}$. $u$ is the input to the hypothalamus (Eq 31) , $b_{i}$ are the secretion parameters, $a_{i}$ the hormone removal rates, $P_{G}$ is the functional mass of the gonadotroph cells in the pituitary that secrete $x_{2},$and $G$ is the functional mass of the gonadal cells that secrete $x_{3}$. The equations model the negative feedback of $x_{3}$on the secretion of x_1_ and x_2_ by means of 1/$x_{3}$ terms, which approximate Michaelis-Menten forms 1/(𝑘+$x_{3}$) in the limit where hormones exceed their Michaelis constant $x_{3}$≫𝑘. Two equations describe the effect of the hormones on the gland masses. GnRH activates $P_{G}$ proliferation (Sakai et al., 1998), and LH/FSH activate G proliferation (Allen et al., 2004; Ma et a., 2004), parameters available at table S3. Simulation was done as for the HPA axis described in Methods.

##### **Growth axis (HP-Liver)**

The growth axis hormone cascade (hypothalamic-pituitary-liver axis) with the feedback by IGF1 on upstream hormones, together with gland mass changes, is described by:

(37) $\frac{dx_{1}}{dt}=\frac{b_{1}u}{x_{3}}-a_{1}x_{1}$
(38) $\frac{dx_{2}}{dt}=\frac{b_{2}P_{S}x_{1}}{x_{3}}-a_{2}x_{2}$
(39) $\frac{dx_{3}}{dt}=b_{3}Lx_{2}-a_{3}x_{3}$
(40) $\frac{dP_{S}}{dt}=P_{S}(b_{S}x_{1}-a_{S}\boldsymbol{)}$
(41) $\frac{dL}{dt}=L(b_{L}x_{2}-a_{L})$

Where concentration of GHRH is $x_{1}$, GH is $x_{2}$ and IGF1 is $x_{3}$. $u$ is the seasonal input to the hypothalamus (Eq 31), $b_{i}$ are the secretion parameters, $a_{i}$ the hormone removal rates, $P_{S}$ is the functional mass of the somatotroph cells in the pituitary that secrete GH, and $L$ is the functional mass of the IGF1-secreting cells in the liver. Negative feedback by IGF1 on secretion of its upstream hormones is modeled as above using 1/$x_{3}$ terms, which approximate Michaelis-Menten terms 1/(𝑘+𝑥3) in the limit where hormones exceed their Michaelis constant 𝑥3≫𝑘. Two equations describe the effect of the hormones on the gland sizes. GHRH activates $P_{S}$ proliferation (Mayo et al., 2000; Solloso et al., 2008), and GH activates L proliferation (Krupczak-Hollis et al., 2003; Pennisi et al., 2004), with similar logic to the HPA axis. Unlike the HPA axis, the timescale of the two tissues is very different. Liver hepatocytes have a half-life of about 300 days or longer (MacDonald, 1961), which is at least ten times slower than the estimated half-life of the pituitary somatotroph cells. This affects the hormone phases: instead of a summer peak for GH, the model shows a spring peak (together with the effector hormone IGF1), as observed in the Clalit data in Fig 1. Seasonality was simulated in a similar way to the HPA, parameters shown in table S3.

##### **HPT axis**

The hypothalamic-pituitary-thyroid axis, together with gland mass changes, is described by

(42) $\frac{dx_{1}}{dt}=\frac{b_{1}u_{T}}{x_{3}}-a_{1}x_{1}$
(43) $\frac{dx_{2}}{dt}=\frac{b_{2}P_{T}x_{1}}{x_{3}}-a_{2}x_{2}$
(44) $\frac{dx_{4}}{dt}=b_{3}Tx_{2}-a_{3}x_{4}$
(45) $\frac{dP_{T}}{dt}=P_{T}(\frac{b_{TT}}{x_{3}}-a_{TT}\boldsymbol{)}$
(46) $\frac{dT}{dt}=T(b_{T}x_{2}-a_{T})$
(47) $x_{3}=\frac{x_{4}}{u_{T}}$

Where concentration of TRH is $x_{1}$, TSH is $x_{2}$ and $T_{3}$ is $x_{3}$and $T_{4}$ is $x_{4}$. $u_{T}$ is the input to the hypothalamus, which we take to be proportional to temperature. We thus use a cosine input function with a peak in August 15, which approximates the mean peak temperature time in Israel. The $b_{i}$ are the secretion parameters, $a_{i}$ the hormone removal rates. The functional mass of the TSH-secreting thyrotrophe cells in the pituitary is $P_{T}$ and the functional mass of thyroxin-secreting thyroid cells (thyrocytes) is $T$. The equations model the negative feedback using 1/$x_{3}$ terms similar to the models above. This assumes for simplicity that $T_{3}$, the main feedback signal, is proportional to $T_{4}$, the main hormone secreted by the thyroid. In reality, we observe a slight seasonal delay between $T_{3}$, and $T_{4}$, suggesting seasonality in the deiodinase system that converts $T_{4}$, to the more active $T_{3}$. To model this, we assume that deiodinases (D) vary inversely with temperature input $u_{T}$, and that $T_{3}=T_{4}D$ (Eq. 47).

To add tissue turnover to the model, two equations describe the effect of the hormones on the functional masses. TSH is the primary growth factor for thyrocytes (Dumont, 1992). Unlike the other pathways, instead of $x_{1}$ being the primary growth factor for the pituitary cells, in this axis it is the effector hormone $x_{3}$ that inhibits growth of the pituitary cells$P_{T}$, as described by the 1/$x_{3}$ term in Eq 45. Moreover, because T4 has a long half-life (~1 week), we model this axis in a slightly different way than the HPA described in Methods. The difference is to account for the fact that the hormones integrate the input u over about a week, whereas in the HPA axis the relevant input is the circadian input during the morning test time. Thus, for the HPT axis, we all numerically solve all 5 equations for the fast and slow timescales together. Parameters are given at table S3.

##### **Prolactin**

Prolactin secretion is inhibited by dopamine and activated by TRH. The Prolactin hormone circuit, together with gland mass change, is described by

(48) $\frac{dx_{1}}{dt}=b_{1}u_{T} x_{2}-a_{1}x_{1}$
(49) $\frac{dx_{2}}{dt}=b_{2}P_{L}/x_{1}-a_{2}x_{2}$
(50) $\frac{dP_{L}}{dt}=P_{L}(b_{P}{/x}_{1}-a_{P})$

Where dopamine is $x_{1}$ and PRL is $x_{2}$.$u_{T}$is the input to the hypothalamus which we take to be the same as in the HPT axis due to the shared TRH input. Additional input which we do not model may arise from photoperiod due to melatonin (Wehr et al., 1993) The parameters, $b_{i}$ are the secretion parameters, $a_{i}$ the hormone removal rate, $P_{L}$ is the total functional mass of the lactotroph cells in the pituitary that secret PRL. The equations model the feedback in which PRL activates dopamine secretion. Tissue turnover is described by Eq 49, where dopamine inhibits tissue growth (Kelly et al., 1997). Parameters available at table S3.

| Pituitary  cell type | Parameter | Value | Gland | Parameter | Value |
| --- | --- | --- | --- | --- | --- |
| Corticotrophs | $a_{C}$ | $\frac{\log\left( 2 \right)}{30\left( days \right)}$ | Adrenal | $a_{A}$ | $\frac{\log\left( 2 \right)}{30\left( days \right)}$ |
| Gonadotrophs | $a_{GT}$ | $\frac{\log\left( 2 \right)}{30\left( days \right)}$ | Gonads | $a_{G}$ | $\frac{\log\left( 2 \right)}{30\left( days \right)}$ |
| Somatotorphs | $a_{S}$ | $\frac{\log\left( 2 \right)}{30\left( days \right)}$ | Liver hepatocytes | $a_{H}$ | $\frac{\log\left( 2 \right)}{300\left( days \right)}$ |
| Thyrotrophs | $a_{TT}$ | $\frac{\log\left( 2 \right)}{30\left( days \right)}$ | Thyroid | $a_{T}$ | $\frac{\log\left( 2 \right)}{30\left( days \right)}$ |
| Lactotrophs | $a_{L}$ | $\frac{\log\left( 2 \right)}{30\left( days \right)}$ | - | - | - |

**Table S3. Parameters table**

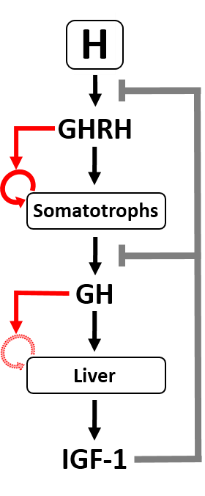

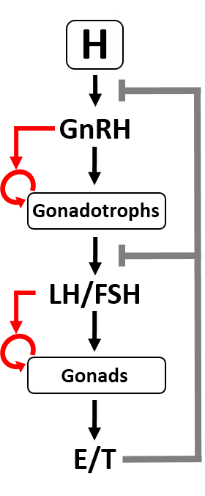

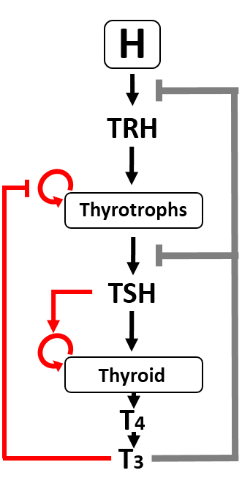

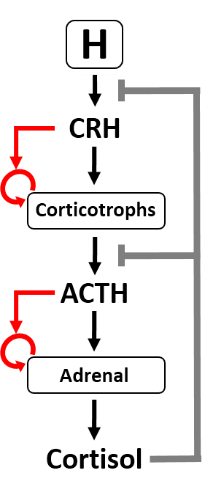

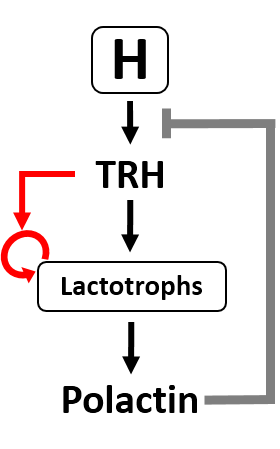

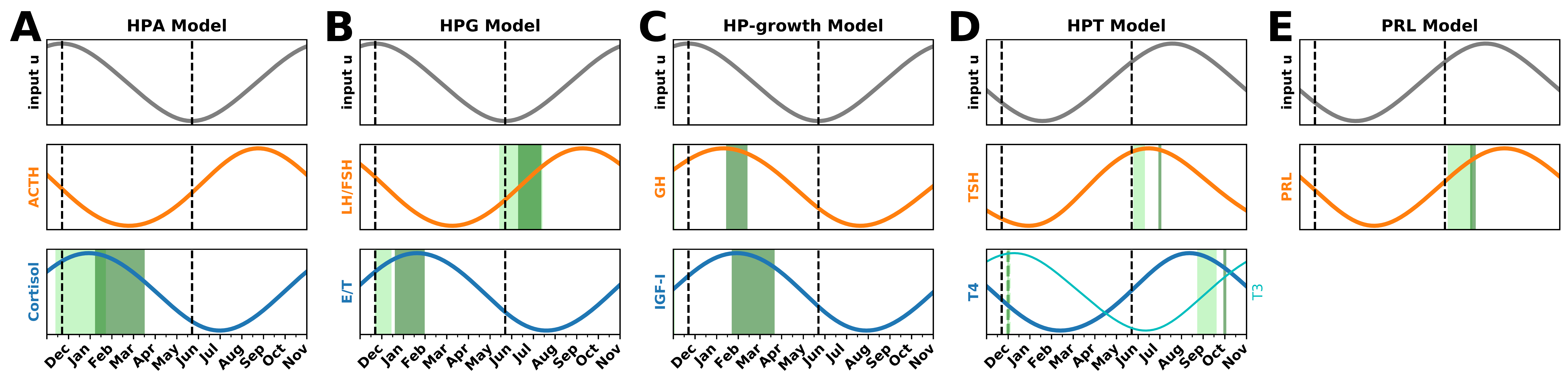

**Fig S11: Models for the hypothalamic-pituitary axes show hormonal phases similar to the observed phases.** Each panel shows the model schematic, with red arrows indicating the effects of hormones on the growth of the corresponding cells. The seasonal input signal is shown, along with the predicted seasonality of the pituitary (orange) and effector (blue) hormones in the linear approximation of the model, valid at small amplitudes. The observed peak phases from the Clalit tests are shown in green (dark green females, light green males), except for ACTH due to the small number of tests. (A) Stress axis (HPA) model, (B) Reproduction axis (HPG) model (C) Growth axis (HP-growth). The hepatocytes unique slow turnover is indicated by the shaded circular arrow (D) Thyroid axis (HPT) model with T4 shown in blue line and T3 in cyan line, (E) Prolactin model (PRL).

#### S10. Alternative mechanisms

We sought to test alternatives to the gland-mass mechanism for hormonal phase shifts. To this, we started with the model without gland dynamics is provided by Eq. 1-3, 6-7. To this model, we added, instead of gland-mass dynamics, four putative alternative biological processes that have a slow timescale, potentially on the order of weeks-months.

(i) The first process is glucocorticoid resistance, where chronically elevated cortisol levels cause weaker feedback from the glucocorticoid receptor GR. To model GR resistance, we added a slow equation that modifies $K_{GR}$, the effective binding coefficient of GR, around its baseline $K_{0}$:
(51) $\frac{dK_{GR}}{dt}={b_{K}K}_{0}x_{3}-a_{K}K_{GR}$
(53) We modeled a putative effect of a seasonal input into GR resistance,

$$K_{GR}=K_{0}u$$

(ii) Another putative alternative process is a seasonal change in cortisol clearance rate. To model this, we added a term $C_{R}$ to the removal rate term in cortisol equation:
(54) $\frac{dx_{3}}{dt}=b_{3}Ax_{2}-\frac{a_{3}x_{3}}{C_{R}}$

Because there is no well-characterized biological process that governs removal on the scale of weeks, we use a phenomenological description in which input $u$ controls cortisol removal directly by setting$C_{R}=u$.

(iii) We also considered a putative model which cortisol reduces is own removal rate by increasing $C_{R}$:
(55) $\frac{dC_{R}}{dt}=b_{R}x_{3}-a_{R}C_{R}$

All of these processes were provided with a month timescale, by setting

$$a_{p}=b_{P}=a_{A}=b_{A}=a_{K}=b_{K}=a_{R}=b_{R}=\frac{\log\left( 2 \right)}{30} days$$

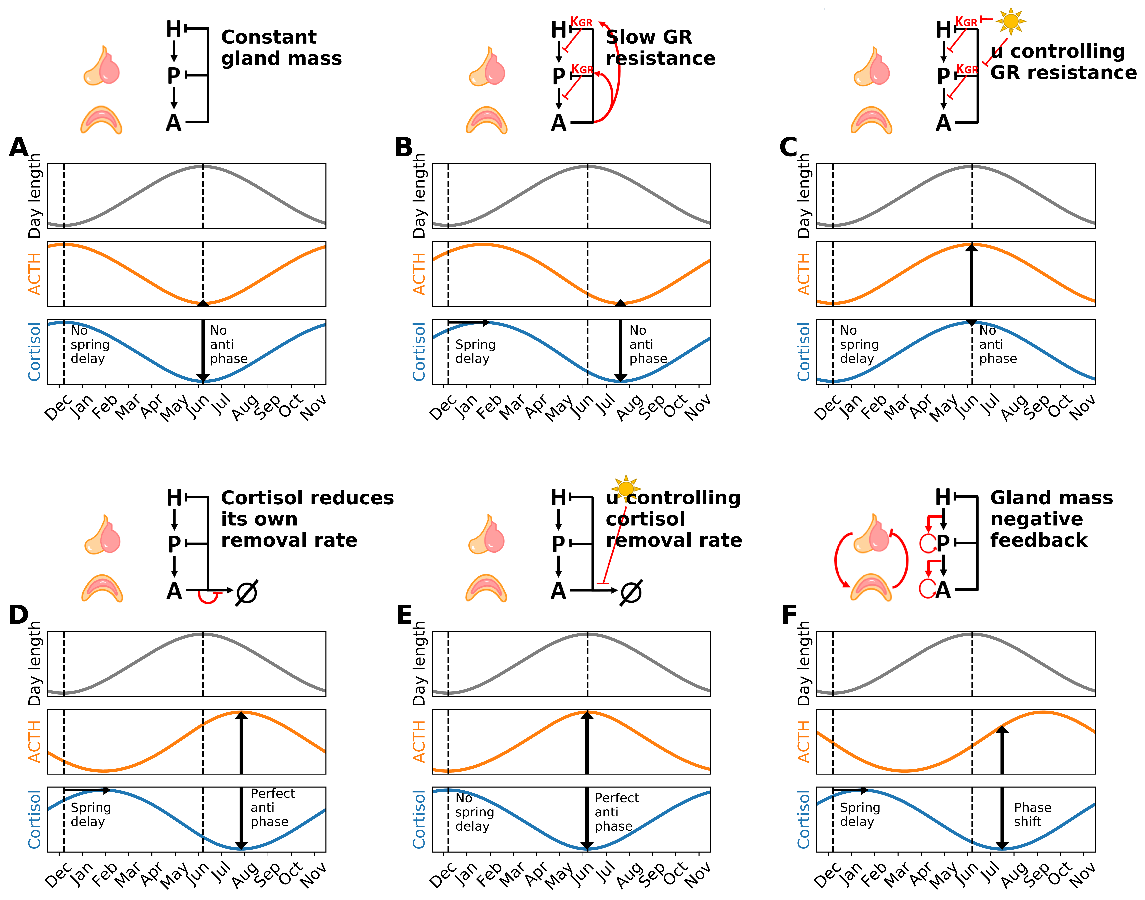
The results are shown in Fig S12. We find that only one of these alternative mechanisms provides both a spring delay to cortisol and an antiphase to ACTH. This is alternative (iv), the reduction of cortisol removal rate by cortisol. It supplies perfect antiphase between the hormones (note that the observed antiphase is only approximate), Fig S12D. The interaction in this mechanism is, however, not supported by experimental data as far as we know. We conclude that out of the models tested here, the gland mass model (Fig S12F) remains the only experimentally supported mechanism for the seasonal phases of the HPA hormones.

**Fig S12: Seasonal hormone dynamics in alternative HPA mechanisms.** (A) The classic HPA model without gland mass changes shows no delay or antiphase in response to photoperiod input (note that HPA input is maximal when photoperiod is minimal). (B) Slow GR resistance caused by high cortisol levels provides a spring delay to both hormones with no antiphase. (C) Putative slow GR resistance controlled directly by photoperiod shows no delay or antiphase. (D) A putative mechanism in which cortisol slows its own clearance provides spring delay and perfect antiphase. (E) A putative mechanism where photoperiod directly affects cortisol clearance shows no spring delay and perfect antiphase. (F) Gland-mass model provides spring delay and approximate antiphase.

#### S11. Reanalysis of previous studies on cortisol seasonality

We used absolute cortisol variation from Clalit rather than percentile changes to compare with the other studies in Fig 4D. We compared to all available cortisol seasonality papers on adults with more than 4 seasonal time points measurements. From each study we obtained the average cortisol values at each time point, and calculated the relative max-min and error by bootstrapping the months. For the Australian study (Hadlow et al., 2018), the relative max-min amplitude was taken from the paper. To estimate the error of the relative max-min amplitude from the Australian study, we used the reported error-bars of summer to winter percentage change (Fig 1 column 2 in Hadlow et al., 2018), and divided this by 2 to estimate the relative max-min error. The computed relative max-min amplitudes and errors are available at https://github.com/alonbar110/Human-hormone-seasonality. See Data and code availability statement”

Studies differed in the way they controlled for circadian effects. In the present study and in the Australian study, cortisol was measured at various times of day in the morning (7:00-12:00). The Swedish study (Persson et al., 2008) used tests at 6:40am. The Whitehall II study (Abbel et al., 2016) used hair measurements which average over 3 cm of hair which corresponds to about 3 months and are thus thought to not be affected by circadian rhythms.

Studies also differed in the filtering of medical conditions and age of participants. The Australian study used a clinical population and removed outlier cortisol measurements and subjects under 18 years of age. The Swedish study used a healthy adult population, with an age range of 32 to 61. The Whitehall study was on UK civil servants aged 59-85, and did not remove subjects due to disease or medication.

These differences, as well as the use of serum, urine, saliva or hair in different studies, are likely to generate different biases in the absolute cortisol levels. Taking relative amplitudes as in Fig 4D probably reduces some of these biases. All of the studies found a cortisol acrophase in winter. Interestingly, a study from Netherlands (Rosmalen et al., 2005) on 1768 primary school children (age 10-12, saliva tests at 7:30 am) showed a summer cortisol peak, perhaps due to proximity to adrenarch.

#### S12. Pituitary volume analysis

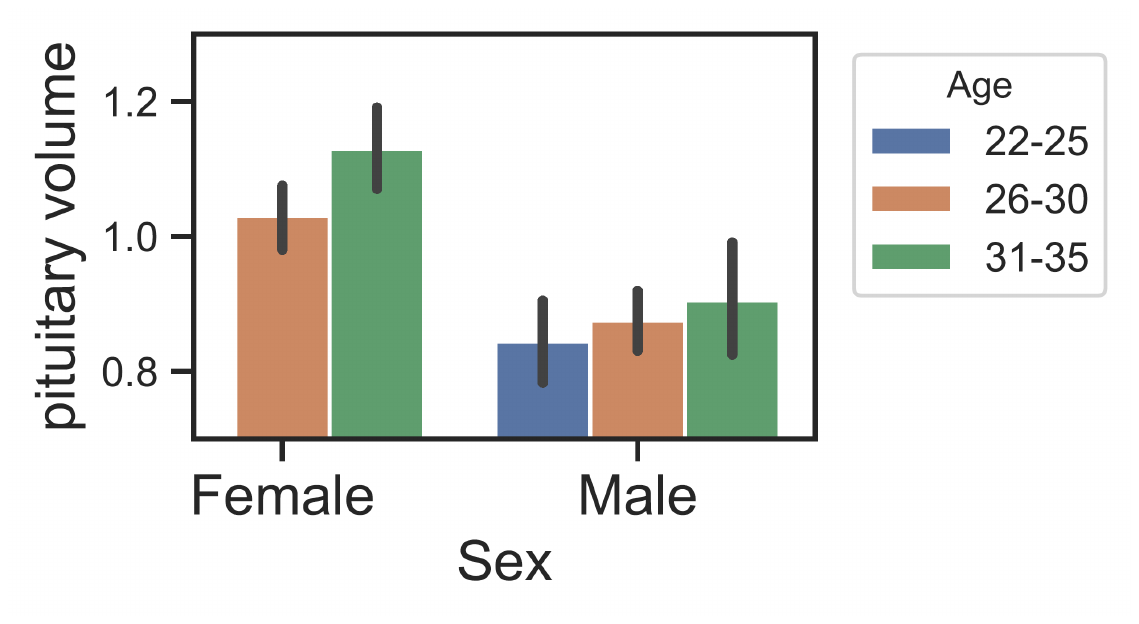
To explore the effects of sex and age on pituitary seasonality, we conducted 2-way ANOVA analysis. This analysis confirmed that there is significant effect of age and sex on pituitary volume (Fig S13). Next, we used linear regression to remove the effects of sex and age. We note that the relative amplitude of pituitary seasonality declined from 6.8% to 5%.

**Fig S13. 2-Way ANOVA analysis for pituitary volume.** Analysis revealed effects for age group and sex on pituitary volume. Full analysis results are shown in table S4.

| **Source** | **SS** | **DF** | **MS** | **F** | **p-value** |
| --- | --- | --- | --- | --- | --- |
| Gender | 1.12 | 1 | 1.12 | 32.1 | <0.001 |
| Age_group | 0.24 | 2 | 0.12 | 3.4 | 0.036 |
| Gender * Age_group | 0.08 | 2 | 0.04 | 1.15 | 0.318 |
| Residual | 5.97 | 171 | 0.04 |  |  |

**Table S4. 2-Way ANOVA analysis summery.**
